# Playbook Workflow Builder: Interactive Construction of Bioinformatics Workflows from a Network of Microservices

**DOI:** 10.1101/2024.06.08.598037

**Authors:** Daniel J.B. Clarke, John Erol Evangelista, Zhuorui Xie, Giacomo B. Marino, Mano R. Maurya, Sumana Srinivasan, Keyang Yu, Varduhi Petrosyan, Matthew E. Roth, Miroslav Milinkov, Charles Hadley King, Jeet Kiran Vora, Jonathon Keeney, Christopher Nemarich, William Khan, Alexander Lachmann, Nasheath Ahmed, Sherry L. Jenkins, Alexandra Agris, Juncheng Pan, Srinivasan Ramachandran, Eoin Fahy, Emmanuel Esquivel, Aleksandar Mihajlovic, Bosko Jevtic, Vuk Milinovic, Sean Kim, Patrick McNeely, Tianyi Wang, Eric Wenger, Miguel A. Brown, Alexander Sickler, Yuankun Zhu, Philip D. Blood, Deanne M. Taylor, Adam C. Resnick, Raja Mazumder, Aleksandar Milosavljevic, Shankar Subramaniam, Avi Ma’ayan

## Abstract

Many biomedical research projects produce large-scale datasets that may serve as resources for the research community for hypothesis generation, facilitating diverse use cases. Towards the goal of developing infrastructure to support the findability, accessibility, interoperability, and reusability (FAIR) of biomedical digital objects and maximally extracting knowledge from data, complex queries that span across data and tools from multiple resources are currently not easily possible. By utilizing existing FAIR application programming interfaces (APIs) that serve knowledge from many repositories and bioinformatics tools, different types of complex queries and workflows can be created by using these APIs together. The Playbook Workflow Builder (PWB) is a web-based platform that facilitates interactive construction of workflows by enabling users to utilize an ever-growing network of input datasets, semantically annotated API endpoints, and data visualization tools contributed by an ecosystem. Via a user-friendly web-based user interface (UI), workflows can be constructed from contributed building-blocks without technical expertise. The output of each step of the workflows are provided in reports containing textual descriptions, as well as interactive and downloadable figures and tables. To demonstrate the ability of the PWB to generate meaningful hypotheses that draw knowledge from across multiple resources, we present several use cases. For example, one of these use cases sieves novel targets for individual cancer patients using data from the GTEx, LINCS, Metabolomics, GlyGen, and the ExRNA Communication Consortium (ERCC) Common Fund (CF) Data Coordination Centers (DCCs). The workflows created with the PWB can be published and repurposed to tackle similar use cases using different inputs. The PWB platform is available from: https://playbook-workflow-builder.cloud/.

## Introduction

High throughput measurements of myriad biomolecules in biological systems have led to generation of large volumes of data creating a paradigm shift in biomedical research. While these large and diverse data spanning a significant dynamic range of length, time scales, and granularity they are highly valuable for deriving new biological knowledge. The ability to discover, access, interoperate, integrate, and analyze these data pose challenges to researchers. As bioinformatics data analyses become increasingly complex and customized, and at the same time more standardized, workflow engines and workflow languages that combine analyses for multiple datasets with a combination of tools are increasing in usage, application, and availability. Hence, a wide array of bioinformatics workflow engines and languages exist (Table S1). Each of these resources has advantages and disadvantages. Broadly, a bioinformatics data analysis workflow platform modularizes data analysis tasks into steps which can be performed in isolation. Capturing dependencies between each step enables stringing them into workflows. Many bioinformatics workflow engines and workflow languages are task-agnostic and operate at the command-line interface (CLI) or within a programming language such as Python. Some of the first generation workflow platforms geared towards bioinformatics were Ruffus [1], Anduril [2,3], Bioconductor workflows [4], and Taverna [5,6]. Ruffus and Anduril are Python libraries that make it easier to combine analysis from multiple tools. Taverna was a larger project that was initially called Taverna Workbench and later Apache Taverna. It could be operated as a desktop application, by CLI, or via a remote execution server and it was coupled with a catalog of workflows called BioCatalogue [7]. With the arrival of the cloud and due to rapid expansion in the availability of bioinformatics tools, the original platforms such as Ruffus and Taverna were superseded with platforms that offered more features and flexibility. These platforms are led by Galaxy [8–11] an internationally large-scale well-funded project that offers many features including a user interface (UI), a library of components, and extensive user training. Alternatives to Galaxy include platforms such as Snakemake [12,13] and NextFlow [14].

These newer platforms rely on community standards that are used to code information about each workflow in a way that enables executing workflows across platforms. The two leading standards are Common Workflow Language (CWL) [15] and Workflow Description Language (WDL) [16]. CWL can be executed by cloud workspaces that implement the Global Alliance for Genomics and Health (GA4GH) Workflow Execution Service (WES) API specification [17], for example, CAVATICA [18] and Terra [19]. Other examples of community standards developed to encode the metadata about workflows include BioCompute Objects, a JavaScript Object Notation (JSON) Schema validatable IEEE standard (IEEE 2791-2020) [20], and WorkflowHub [21] which describes workflows by adopting the Research Object Crate (RO-Crate) standard [22] and leveraging schema entities from BioSchemas [23].

The growing collection of thousands of publicly available bioinformatics tools with APIs also gave rise to another class of system tangentially related to workflow systems. These are federated knowledge graphs (KGs). Examples of such systems include the BioThings Explorer [24] which invokes APIs documented and registered with the SmartAPI registry [25] to dynamically resolve edges between two destination data types. BioThings Explorer is related to NCATS’ Translator project [26,27] which operates similarly but with a UI that looks like a typical search engine. Another class of system tangentially related to workflow engines are UIs that enable the user to upload their data into a cloud environment and then select the tools and other aspects of their desired workflow, and then, once pressing submit, the workflow is executed in the cloud and the results are delivered as a report. For example, BioJupies [28] is a platform for enabling researchers to perform RNA-seq analysis in the cloud. The user can start with a data matrix, or a collection of FASTQ files. After these files are uploaded, the user can pick from a collection of tools that will be executed to produce the data analysis report resembling a Jupyter Notebook [29]. The BioJupies platform was later extended to enable quick analysis of many other data types with Appyters [30]. Appyters are parameterized Jupyter Notebooks converted into full-stack web-based applications. Other similar platforms include GenePattern [31] and iLINCS [32]. Several of the platforms in this category interoperate with other workflow languages, especially with CWL and WDL.

Since its inception in 2004, the US National Institutes of Health (NIH) Common Fund (CF) has funded more than 50 CF programs. CF programs have generated large and diverse datasets with the aim of having these datasets propel biomedical research forward by serving as resources for hypothesis generation and integrative systems level analyses. These datasets include various omics profiling from across thousands of human subjects, cell lines, organoids, and animal models. Each CF program typically has a Data Coordination Center (DCC) that is tasked with managing these datasets and serving them to the community via bioinformatics tools, workflows, and interactive web-based UIs. DCC portals often go beyond serving the raw data from their respective CF program by providing more distilled information and knowledge extracted from such data. To accomplish this, DCCs have in many cases designed tools that enable users to interactively explore datasets via user interfaces as well as via well-documented API. However, enabling knowledge discovery by combining data and tools from multiple CF programs remains both a challenge and an opportunity. To address this challenge, the NIH established the Common Fund Data Ecosystem (CFDE) consortium. In its first phase, the CFDE consortium established common data elements and harmonized descriptors for biological and biomedical entities such as genes, tissues, drugs, and diseases across centers and programs [33]. These harmonized descriptors can be used to describe raw files produced by each KF program but fall short of directly enabling cross-program hypotheses generation.

Here we demonstrate how by leveraging data, tools, and well-documented FAIR representational state transfer (REST) APIs from multiple CF programs, and other sources, we constructed a visual user-friendly web-based workflow construction platform called the Playbook Workflow Builder (PWB). In contrast with other interactive workflow platforms, PWB requires stricter annotations and specifications of workflow components that we term metanodes. Such extensive descriptions of metanodes provide users with a richer, more focused, user experience that enables advanced and complex data analyses, data harmonization, and data integration. Specifically, users can interactively and visually create workflows by exploring all possible available options at each workflow construction step. The PWB consists of a growing network of connected metanodes, and it is demonstrated via 12 published workflows that are served as reports that resemble parameterized publications.

## Methods

To dynamically develop workflows that draw knowledge from across CF programs and other key bioinformatics tools and databases, we synthesize information stored in multiple CF DCCs to accumulate evidence about a specific hypothesis. To achieve this, we integrated several CF DCC tools and databases accessible via well-documented APIs into an integrative network of DCC microservices. odes in the network represent semantic types, for example, gene sets, gene expression signatures, diseases, metabolites, glycans, and drugs. Edges in the network represent transformations, or operations, performed by various tools on these semantic types, for example, enrichment analysis applied to a set of genes or a set of metabolites, principal component analysis (PCA) applied to a data matrix, or a PubMed search applied to a search term that describes a disease. Nodes and edges are characterized in a strict type-safe manner forming a programmatically defined data structure we term a knowledge resolution graph (KRG). In contrast to a KG, a KRG encodes the capacity of obtaining knowledge by means of some computational or manual process; instead of subjects connected via predicates, as in a KG, a KRG has functions connected via common data types. Knowledge obtained from one tool or database may be augmented, compared, or supplemented with knowledge from another. The KRG can be used to help users find compatible processes with instantiated knowledge at any step in a chain of steps, forming a complete workflow. The tools collected are largely REST API-driven microservices providing complementary interoperability across CF DCCs and other relevant biomedical databases and tools. The APIs are documented with OpenAPI [34] and deposited into the SmartAPI [25] repository. Such compliance with these standards eases implementation.

The assembled metanodes are then used to facilitate a collection of use cases and use case templates. Use case templates are defined as workflows with the same structural components but with application to different data instances. For example, gathering information about a gene or a variant from several CF DCCs databases can be done for a single gene, but also as a template that supports the querying of other genes by changing the input query. The collected use cases are geared toward accumulation of evidence from transcriptomics, metabolomics, glycomics, proteomics, epigenomics, genomics, imaging, and other assay types. The workflows that are generated for realizing these use cases are reusable and extendible. To enable access to the system, a user-friendly interface (UI) was developed. The UI is geared to experimental biologists with no programming background. The PWB system is set up in a way that other developers can contribute to the system, and/or reuse components of the PWB for enhancing their own web portals and bioinformatics data analysis workflows. Metanodes are accessible via a uniform REST API that supports multi-step workflow executions via CWL. Thus, the KRG graph can be queried programmatically.

### Metanode Specifications

Metanodes are specified with TypeScript. The specification captures common identifiable metadata elements about each component. These include human readable labels, descriptions, icons, authors, license, and versioning information. The specification then couples these semantics with type-safe implementations which inherit types from dependent components. The set of components is compile-time checked and can be queried and operated-on at runtime in a type-safe manner through runtime-based type checking. A metanode can be a *data type*, a *resolver*, or a *view* (Fig. 1). A *view* function renders the visualization of an instance of the type of interface. A *resolver* function accepts one or many *data types* as inputs and produces a single *data type* as an output. A *prompt* is a React component that can accept input *data types* to facilitate decisions made by the user for transforming the inputs into a single output *data type*, for example, selecting a gene from a list, or submitting a gene set for enrichment analysis. With these three metanode types, we can construct workflows. A *prompt* with no inputs can inject an initial instance of a *data type* object, and that instance can be used as an input argument to compatible *resolvers* or *prompts* to yield other *data type* instances, or figures, tables, and charts. Metanodes also specify parts of a story. This is a parameterized sentence about what that component is doing as it might appear in the methods section of a research paper. These sentences are stacked together to construct a human-readable description of the entire workflow. The paragraph can be further reorganized and copyedited using an LLM like GPT-4 [35].

**Fig. 1.**
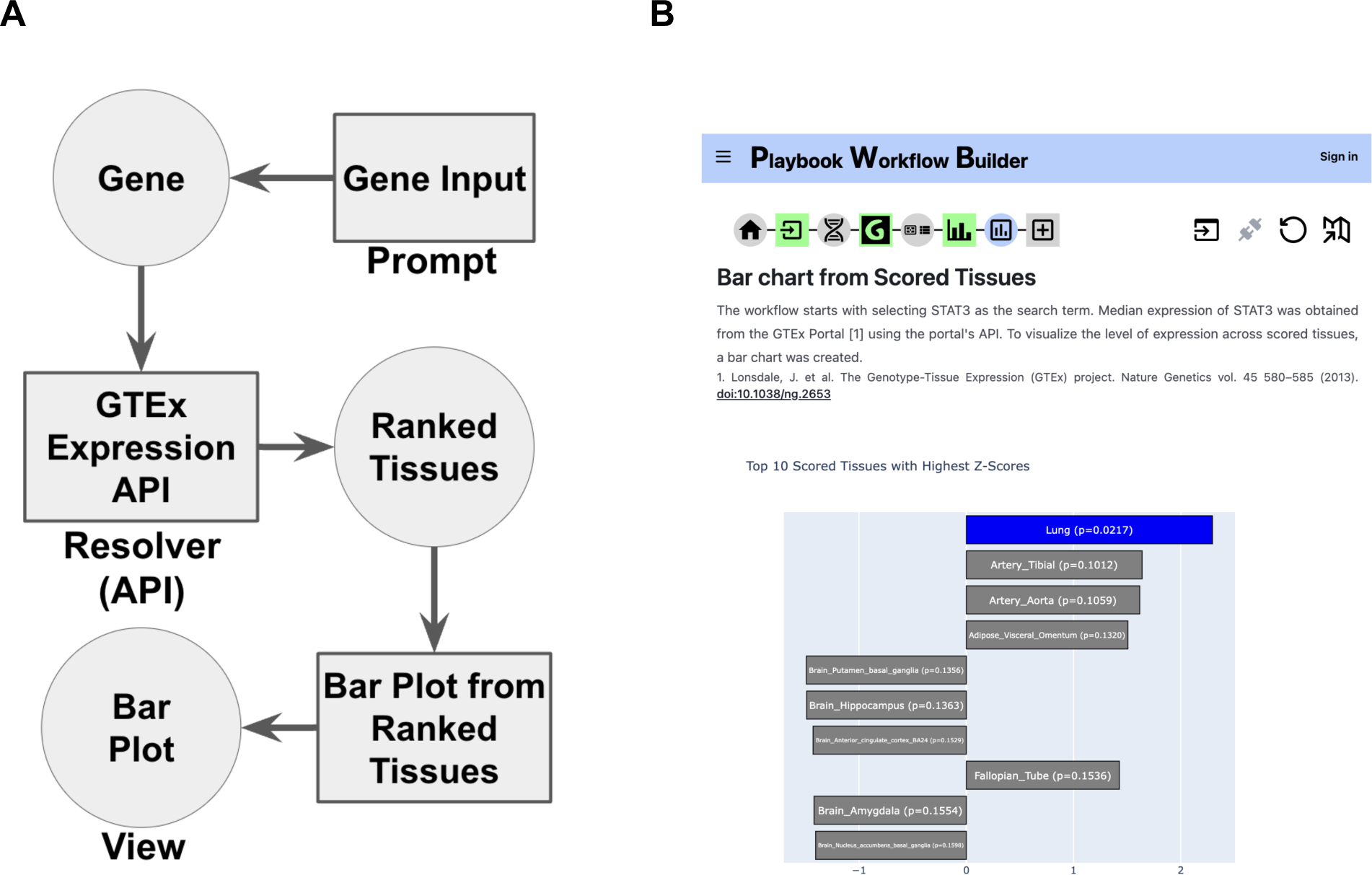
The different PWB metanode types and how they are strung together to form workflows. **A.** In this example, the prompt type of metanode takes a gene as the input; then the resolver metanode uses the GTEx API to obtain the expression of the input gene from across human tissues. Finally, a view metanode visualizes the contents returned from the API as a bar chart. **B.** Screenshot from the executed workflow in the PWB platform.

### Knowledge Resolution Graph (KRG)

Because of the metanode specification, PWB metanodes can be developed, tested, and operated independently from the PWB codebase. All the implemented metanodes are collected and assembled into a unified KRG database (Fig. 2). The PWB system queries and utilizes this database to construct the data-driven UI (Fig. 3). As such, the PWB web-based application is a product of the contents of the KRG database, and thus, extending the functionality of the PWB web-based application only requires creating and registering additional metanodes. By modularizing the PWB processes we can mix, match, and stack PWB metanodes to construct parameterizable workflows. PWB metanodes and workflows have consistent interfaces and can thus be exposed in consistent ways such as over API, in CWL workflows, or through visual interfaces.

**Fig. 2.**
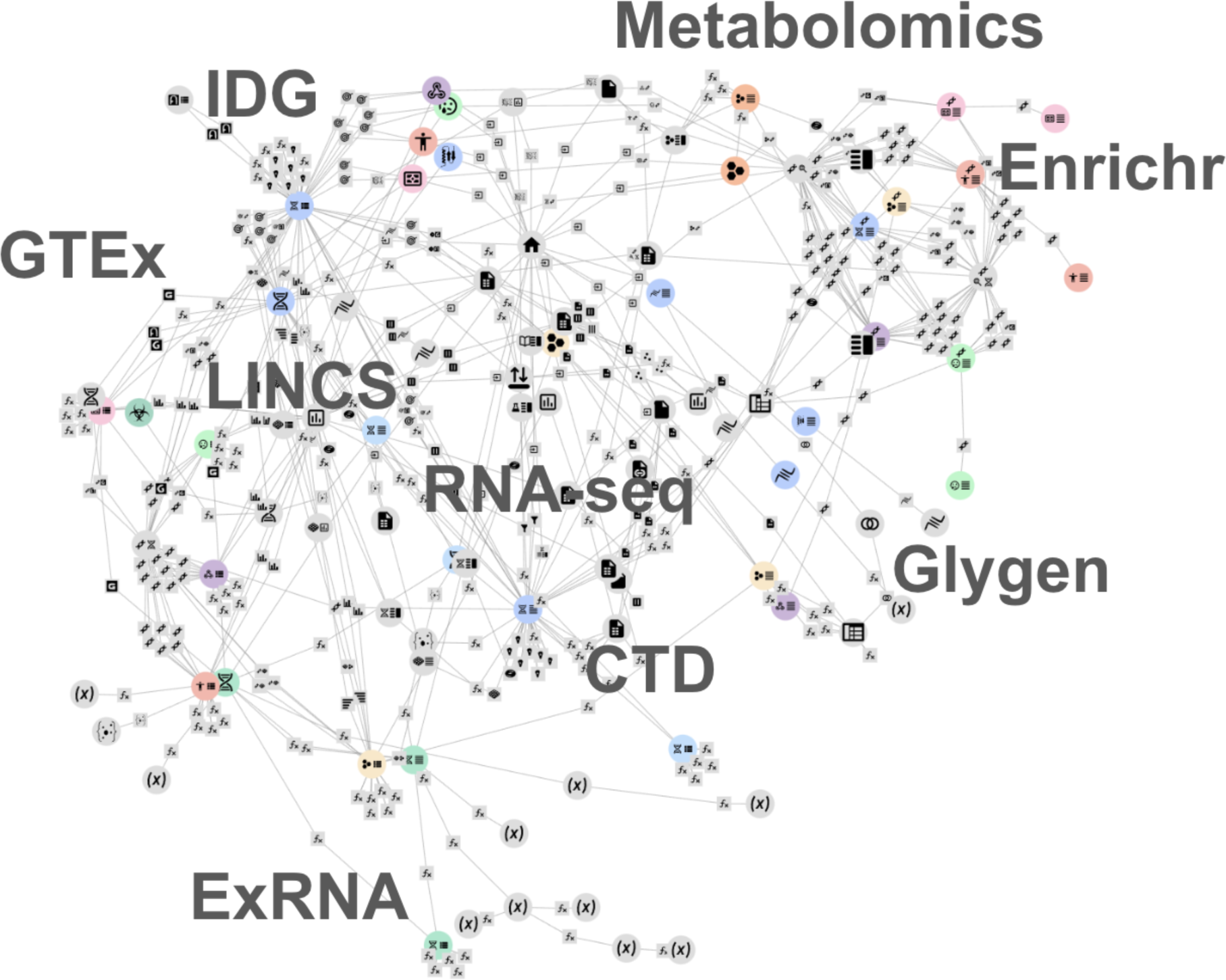
Network visualization of the PWB knowledge resolution graph (KRG). The network of connected metanode is interactive and can be explored from the user interface.

**Fig. 3.**
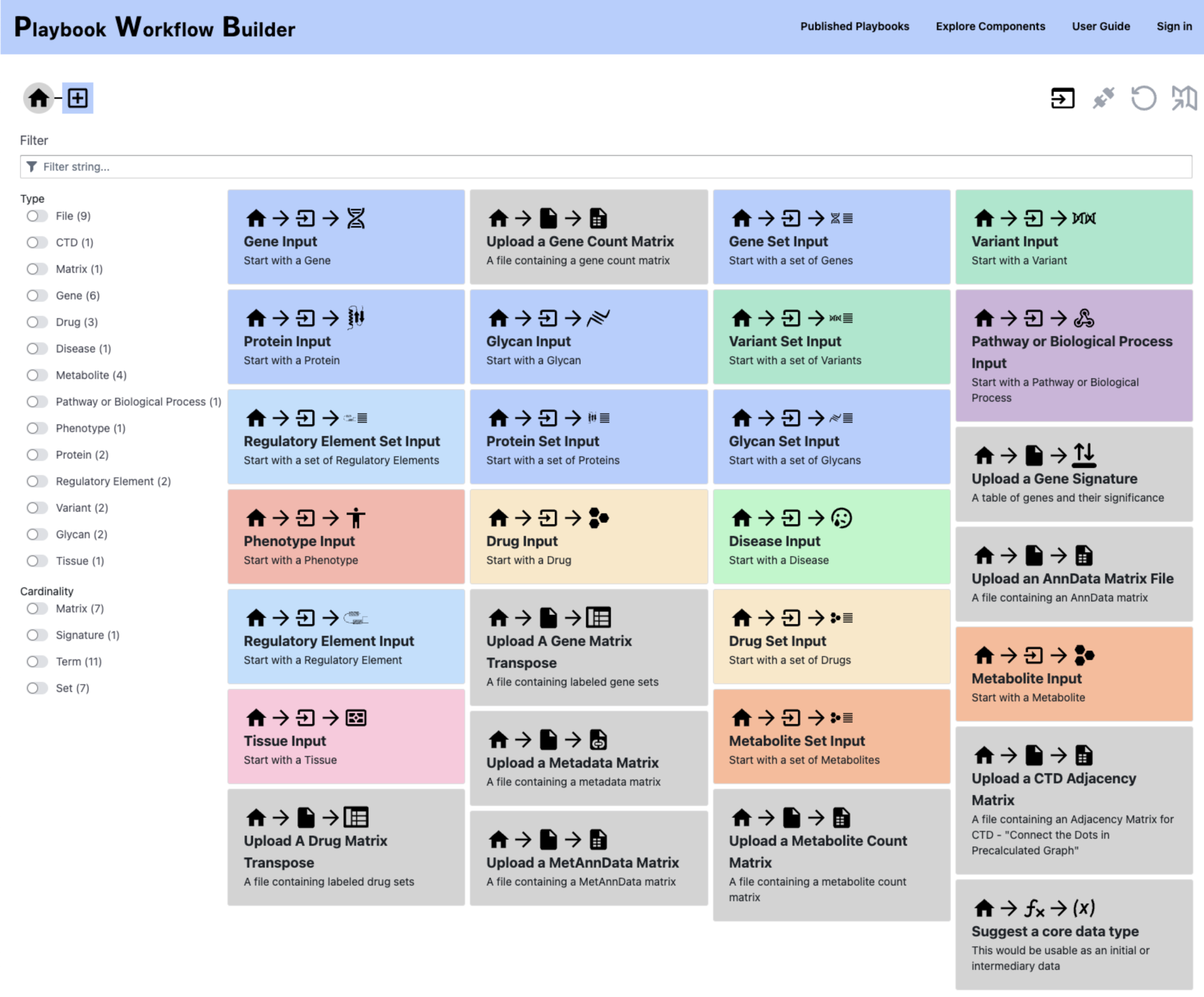
The landing page of the PWB UI provides access to a collection of prompt metanodes to begin constructing workflows.

### Fully Persistent Process Resolution Graph (FPPRG)

While the KRG can be used to construct arbitrary workflow templates, a workflow is an instance of that template operating on a unique dataset that has the same structure but different contents. To store data from a workflow, an additional database is established. This additional database stores the data that flows through workflows. As such, the database ensures collision-free updates and a self-deduplication. Another feature of this additional database is the decoupling of workflow templates from the actual data that flows through those workflows, providing further deduplication. Workflows are stored across four independent tables (Fig. S1). The first table is a dependency graph of each constructed step of a workflow. This information is stored in a record called a *Process*. This record is tightly coupled with the *Component*, it stores the *Component* ID, a JSON object for *Prompt* configuration, and back references to any other *Process* whose output is used by this record. The second table is a fully persistent list (FPL). It stores sequential order of a workflow through a linked list. A singular list can be resolved with the ID of the last element of the list, and each intermediary state has a unique ID. Importantly, elements of the lists need only be stored once even if used in multiple lists. The third table is a *Result* record. It has a one-to-one relationship with a *Process* record and is constructed by performing the execution using the function from the *Component* type referenced in the *Prompt*. Finally, the 4th table is a *Data* record. This table contains arbitrary JSON Binary Large Objects (BLOBs), used to store data in the *Process* and *Result* tables. All IDs are created by hashing the content of the record. A unique series of user steps can be stored and accessed by a single ID through the FPL, while the dependency graph ensures deduplication of the workflows regardless of order. Finally, the actual results of any given workflow step are stored. Requests for the output of any *Process* are sent to a queue of workers if the *Result* does not already exist. Hence, steps are executed simultaneously if there are enough workers, and equivalent execution results are deduplicated. Altering an earlier step in a workflow can be done with a git-style rebase. A new FPL and dependency graph starting from the parent of the modified node are created and expanded to the previous tail. *Result* records would then be computed as required to obtain the new output for the entire workflow.

### Developing the PWB Website

The PWB website is developed in TypeScript with NextJS, a full-stack framework that uses React and offers isomorphic server-side and client-side rendering. TailwindCSS-based DaisyUI and BlueprintJS are used for styling the site and data tables. NextAuth.JS is used for managing user accounts via ORCID or e-mail. The FPPRG which stores workflow executions can operate entirely in memory or with a PostgreSQL database in a production setting. Workers run in the main process or execute independently on different machines. Message passing is achieved through PostgreSQL’s listen/notify feature. The website’s navigation and metanode rendering are driven by queries to the in-memory KRG over REST API or WebSocket. The UI is decoupled from the metanodes facilitating the independent development of the website and the metanodes. This also means that a completely new set of metanodes can be used for a platform with a different focus. All metanode TypeScript, Python, and other dependencies are assembled and installed into a single Docker container. This container is used to run the PWB workers. A smaller Docker container with only JavaScript dependencies runs the UI.

### Cloud Agnostic File Storage

A Python library was developed to help with managing files in a storage system that is agnostic to the cloud provider. All files uploaded to the PWB are stored and accessed using an abstract layer provided by this library. In development, files are stored on the local disk, while in production, the files are stored in an S3 bucket. Alternatively, users can have their files in a CAVATICA workspace [18] when CAVATICA sessions are established. Once uploaded, files are stored by their sha-256 checksum which provides content-based addressing for deduplication. An entry is added to the database and is associated with the user who uploaded the file. These records receive universally unique identifiers (UUID) and are served by the PWB platform using GA4GH’s Data Repository Service (DRS) protocol [36]. Files on the platform are then treated as DRS URIs which can be resolved anywhere in the system. Files can also be provided to the platform directly from external DRS hosting platforms. Functional helpers are available to obtain the contents of the DRS files/bundles or for uploading new files from within PWB metanodes.

### Workflow Format Translations

The FPPRG format encodes the workflows along with the data that flows through these workflows. The steps of the workflow are encoded in the KRG where metadata about the steps can be resolved. These can thus be translated to other community developed workflow description formats for the purpose of interoperability with other platforms. Hence, the PWB platform provides users with the ability to export constructed workflow into several workflow specification standards.

#### BioCompute Objects

There has been a need in bioinformatics for establishing better conceptual descriptions of workflows [37]. For example, a bioinformatician may wish to reproduce a pipeline using tools or platforms that they are familiar with and which they trust. Workflow languages – machine-readable files that confer portability of execution – are usually insufficient when context and a conceptual underpinning is needed. For this, the BioCompute objects standard was developed [20] IEEE 2791-2020. BioCompute is a rigorously defined standard for bioinformatics analysis workflow documentation that is flexible enough to accommodate any pipeline, but rigid enough to define a structure for computable metadata to annotate workflows. There is an ecosystem of tools for working with the BioCompute standard, including large cloud genomics platforms like Seven Bridges Genomics, CAVATICA, and DNAnexus. The BioCompute Portal is part of this ecosystem, and acts as a repository of published BioCompute Objects (BCOs), as well as a place to manually build BCOs, and has been used in two published examples [38,39]. On the PWB, A BCO can be constructed from any given FPPRG, containing full provenance about the workflow including the individual steps and authorship information. These serialized BCO specifications can be downloaded or directly sent to the BioCompute Portal via API where they can be inspected, modified with additional annotation, or extended to other schemas, and ultimately published.

#### Common Workflow Language (CWL)

Common Workflow Language (CWL) is an open standard for describing how to run command line tools and connect them to create workflows [15]. A command line interface (CLI) was developed from the KRG to invoke any *Process* metanode, providing inputs in JSON-serialized files, and writing the output to a JSON-serialized file. Using this CLI, a CWL CommandLineTool specification can be constructed out of any *Process* metanode, and a CWL workflow specification and input variables file can be constructed out of a FPPRG. All *Prompt* data which would have been captured by the user via a UI are instead specified in the input variable file. Every step of the workflow is exposed as an output. Hence, the PWB platform metanodes are fully compatible with CWL, and CWL workflows can be exported from the PWB interface.

#### Research Object Crate (RO-Crate)

RO-Crate is a community-based specification for research data packaging of Research Objects (RO) with rich metadata, based on open standards and vocabularies including JSON Linked Data (JSON-LD) and schema.org [40]. Adopting a similar structure to describing workflows as WorkflowHub [41], an RO-Crate can be created from an FPPRG. The RO-Crate can then be used for registering PWB workflows in WorkflowHub and for minting citable Digital Object Identifiers (DOIs) for published workflows.

### Constructing Workflows from Prompts with GPT

The user interface of the PWB facilitates construction of workflows by presenting to the user all possible next steps compatible with the current step. This functionality is also presented as a prompt to a large language model (LLM) Assistant, such as generative pre-trained transformer (GPT), capable of making decisions about the best next step to take when presented with a prompt from the user. Using a few-shot prompt, we direct the assistant to choose from a set of possible next steps based on user messages. We accept single suggestions automatically and present multiple suggestions to the user. Selected suggestions are included in an incrementally constructed workflow and rendered in a chat box-style interface along with LLM Assistant messages. Because we use the assistant to only help build a PWB workflow based on the constrained KRG, the risk of hallucination is mitigated. In the worst case, users receive a self-documented workflow that may perform an analysis that is not intended. By collecting feedback from the user in the form of thumbs up or thumbs down, we can fine-tune the model in the future to build more accurate workflows based on user prompts.

## Results

### Implemented Metanodes

The PWB platform provides users with the ability to perform a wide variety of analyses powered by the network of metanodes. These metanodes are used as steps in workflows. So far, we have developed 561 such metanodes (Table S2). Below we describe some of the currently implemented metanodes.

#### RNA-Seq Data Analysis and Visualization

Beginning from a user-uploaded count matrix of gene expression, where each row represents a gene, and each column is a sample with associated metadata, data is uploaded to the PWB and encoded with AnnData [42]. From the gene expression matrix, several metanodes enable different normalization and data visualizations via PCA [43], UMAP [44], or t-SNE [45]. These metanodes are supported by the Scanpy Python package [46]. The data matrix can also be used for computing differential expression to produce gene expression signatures. Differential expression analysis is performed by methods such as the Characteristic Direction [47], limma-voom [48,49], or DESeq2 [50]. Differentially expressed genes can be used as input for downstream analysis such as enrichment analysis.

#### Enrichment Analysis

Enrichment Analysis can be performed within the PWB using the Enrichr API [51]. The gene sets *data type* in the PWB can be enriched against many gene set libraries stored within Enrichr. For example, the GTEx [52] and ARCHS4 [53] libraries can be used to obtain a prioritized ranking of tissues, KEGG [54] and WikiPathways [55] libraries can be used to prioritize most relevant pathways. Enrichr also provides an API to search for metadata terms across the gene set libraries. For example, a disease term search can be used to construct a consensus gene set from GEO disease signatures [56]. Another way to build such gene sets is through literature search based on term-gene co-mentions in publications. This functionality is supported by PWB components that utilize the Geneshot API [57].

#### Gene Set Manipulation

The gene matrix transpose (GMT) file format is commonly used to serialize gene set libraries. GMT files contain lists of terms followed by sets of genes for each term. GMT files are loaded and manipulated in the PWB. A common way to interrogate the overlap between several gene sets through UpSet plots [58] or with a SuperVenn diagram [59]. The PWB has a metanode to display interactive versions of both, enabling users to inspect regions of overlapping and unique genes and gene sets. Additionally, several operations were implemented to transform *data types* from one to another. For example, turning ranked lists of genes into gene sets by choosing a cutoff, turning multiple gene sets into a GMT, or collapsing a GMT into a single gene set by applying a consensus or a union set operation.

#### Healthy Human Tissue Expression Atlases

The NIH GTEx CF program has profiled gene expression data from healthy human tissues [52]. The GTEx API can be used to find median tissue expression levels for all human genes for each one of 54 profiled tissues. Similarly, the ARCHS4 resource [53] was created by uniformly aligning approximately 2 million publicly available RNA-seq samples collected from human and mouse. The ARCHS4 API can also be used to find median tissue expression across over 200 tissues and cell types. The PWB enables users to obtain summary statistics from these APIs which can be visualized as bar graphs. It is also possible to use these data resources as a baseline to identify novel drug targets. For example, gene expression data collected by RNA-seq from tumor samples, can be compared to all normal tissue to identify genes that are only highly expressed in the tumor using the TargetRanger API [60].

#### Metanodes Created from LINCS Resources

The Library of Integrated Network-Based Cellular Signatures (LINCS) NIH CF program [61] profiled the response of human cells to thousands of chemical and genetic perturbations followed by omics profiling. The PWB provides several components related to prioritizing drugs and preclinical small molecules for targeting individual genes and gene expression signatures. Several metanodes can be used to perform LINCS L1000 reverse search queries for a given gene, producing visualizations and tables of significant LINCS L1000 chemical perturbagen signatures which maximally increase or decrease the expression level of the single human gene. Similar metanodes were implemented to provide search against the L1000 CRISPR KO signatures. Other metanodes enable users to query the SigCom LINCS database [62] with gene expression signatures. Such signatures may be in the form of a vector of differential gene expression, or up- and down-regulated gene sets. Both types of input signature queries can yield ranked lists of chemical perturbations and CRISPR KOs.

#### Metanodes Created from GlyGen Resources

GlyGen is an international initiative funded by the NIH to promote research about glycoscience [63]. The GlyGen consortium developed a web-based portal that brings together glycan and protein specific data from major resources such as UniProt [64], GlyConnect [65], Protein Data Bank (PDB) [66], UnicarbKB [67], ChEBI [68] and PubChem [69] and other resources [70]. These datasets are presented to users through a standardized data model [71] via the GlyGen data portal (https://data.glygen.org). The GlyGen API endpoints (https://api.glygen.org) facilitate the same functionality provided by the user interface, providing the PWB with several GlyGen metanodes that can be integrated in various workflows. The GlyGen metanodes also support data visualization and kinase enrichment analysis. Furthermore, the GlyGen metanodes operate several core data types such as, glycans, proteins, and glycoproteins. For other glycoconjugate species, such as glycolipids, GlyGen metanodes implement the passthrough search APIs to the GlySpace alliance [72] and other resources. In addition, uploaded mass spectrographic glycan files are analyzed with various GlyGen specific metanodes, and then knowledge is extended with other PWB metanodes.

#### Metanodes Created from Metabolomics Resources

The Metabolomics Workbench (MW) is another resource supported by the NIH CF [73]. MW contributed several metanodes to the PWB including those from the bioinformatics tools MetGENE [74], MetENP [75], and a gene ID conversion tool. These tools, originally designed to be stand-alone web applications, provide REST APIs to obtain relevant information for analyses related to profiled metabolites within the PWB. MetGENE is a hierarchical, knowledge-driven tool designed for gene-centered information retrieval. By entering a single gene, or a set of genes, users can access information related to the gene such as pathways, reactions, metabolites, and studies from metabolomics in MW. To refine searches, MetGENE incorporates filtering options based on organism, tissue or anatomy, and disease or phenotype. This feature provides tailored and context-specific search experience. Several metanodes using MetGENE are implemented that take as input either a gene, or a gene set, for downstream analyses. The relevant functionality from MetENP is provided via a REST API called MetNet. Briefly, given a list of metabolites, e.g., metabolites with significant change between two conditions such as disease/normal or treatment/control in a metabolomics study obtained by using MetENP or another tool, a researcher may want to find what are the pathways and functions affected. MetENP/MetNet facilitates metabolite name harmonization using RefMet [76], metabolite class enrichment, metabolic pathway enrichment and visualization, and identification of reactions related to the given metabolites and genes coding for enzymes catalyzing these reactions. In MetNet, the list of these genes can be used to develop their protein-protein interaction (PPI) subnetwork using the STRING database APIs [77]. Each of these metanodes have an associated table that renders the information obtained from the API.

#### The Connect the Dots (CTD) Metanode

The Connect the Dots (CTD) metanode takes as input a set of genes or proteins and identifies a subset of genes or proteins that are highly connected within either knowledge graphs or networks derived from gene expression, metabolomic or other omic datasets [78]. CTD algorithm has previously discovered multi-gene biomarkers of drug response to breast cancer therapies based on mouse PDX models [79], and metabolomic signatures of rare inborn errors of metabolism [78,80]. While CTD has been previously deployed as independent R and python packages (https://github.com/BRL-BCM/CTD), its deployment on the Playbook will allow for its use by a wider scientific audience. The CTD workflow starts with an input set of genes. The user then has the option of identifying significant connections within this set in the STRING protein-protein interaction network [77], WikiPathways [55], or a network derived from user-supplied data. The networks represented as weighted graphs, can be derived from expression data, proteomic data, metabolomics, or any other normalized omic dataset. This allows for users to identify highly connected sets of genes within their specific disease, treatment, or condition of interest. Given a weighted graph and a set of graph nodes as an input, CTD identifies significant highly connected subsets. An optional “guilt by association” feature identifies neighboring nodes using probability diffusion. CTD also returns a visual display of the nodes and connections.

#### Metanodes Created from ERCC Resources

The ExRNA Communication Consortium (ERCC) Common Fund (CF) Data Coordination Center created a framework and toolset for FAIR data, information, and knowledge that delineate the regulatory relationship between genes, regulatory elements, and variants, and made them available to PWB via metanodes. We have implemented the ClinGen Allele Registry (CAR) and Genomic Location (GL) Registry [81], variant and genomic region on demand naming services, respectively. The CAR canonical identifiers (CAid) or Genomic Location identifiers (GLid) provided are reference genome-agnostic, stable, and globally unique. The ERCC metanodes enable the retrieval and mapping of unique identifiers and other commonly used identifiers, such as dbSNP ids [96], connected through the Allele Registry and GL Registry using the Allele Registry RESTful APIs. Moreover, we have created the CFDE Linked Data Hub (LDH) [82], a graph-based database, to extract and link tissue and cell type-specific regulatory information from SCREEN [83], GTEx [52], and other CF projects, including Roadmap Epigenome [84] and EN-TEx [85]. Each excerpt on the CFDE LDH is created in a machine-readable format and contains a link to the original data source for provenance tracking. The CFDE LDH RESTful APIs provide read and write capabilities for both accessing and contributing gene regulatory information. This enables the CFDE LDH to connect more than 800 million regulatory data and information documents, which can be quickly retrieved by PWB through the API endpoints given any variant, regulatory region, or gene as input.

### The Book of Use Cases

The PWB currently contains a collection of implemented and published workflows. These workflows were first designed by drawing the workflow as workflow diagrams in a Google Slideshow (Fig. S2). Each slide represents a unique workflow contributed by members of the project. In these diagrams each node represents a metanode. The slide representing a workflow also lists the name of the workflow and the resources used to obtain the data needed to run the workflow. The color of each metanode was used to track the status of implementation of the metanode and the overall workflow. The plots were used as a guide to capture ideas about potential workflow. Thus, not all these use cases are fully implemented and in some cases that actual implemented workflow does not match exactly the diagram that is associated with it.

### Use Case Workflow Templates and Workflow Instances

The PWB implemented published workflows are listed on a dedicated area on the PWB site (Fig. 4). Each published workflow has a title, a short description, a description of the inputs and outputs, the data resources used, the authors, version, license, the date of publication, and a button to launch the workflow. Since each workflow is parameterized, we consider these workflows as playbooks. These playbooks can be executed with different inputs to produce a new workflow. Below we describe several selected published PWB workflows in detail.

**Fig. 4.**
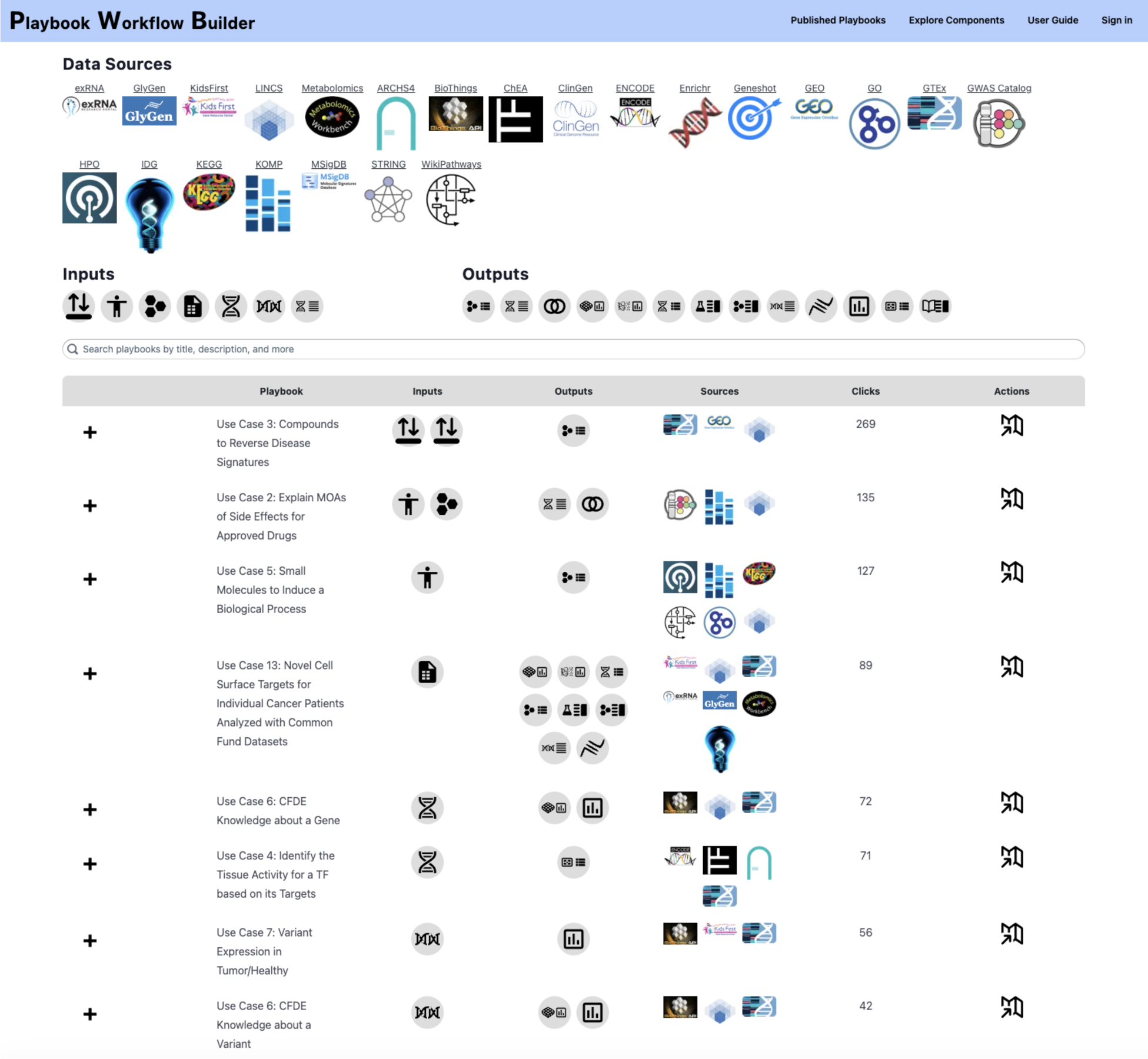
Published workflows are curated workflows that are listed on a dedicated page that lists these in a table. Each workflow entry can be expanded to obtain more information about the workflow and launch the workflow within the PWB platform in report mode.

#### Use Case 13: Cell Surface Targets for Individual Cancer Patients Analyzed with Common Fund Datasets

The input to this workflow is a data matrix of gene expression that was collected from a pediatric tumor from the Kids First CF program [18]. The RNA-seq samples are the columns of the matrix, and the rows are the raw expression gene counts for all human coding genes. This data matrix is fed into TargetRanger [60] to screen for targets that are highly expressed in the tumor but lowly expressed across most healthy human tissues based on gene expression data collected from postmortem patients with RNA-seq by the GTEx CF program [52]. Based on this analysis, the gene Insulin-like growth factor II m-RNA-binding protein 3 (IMP3) was selected because it was the top candidate returned from the TargetRanger analysis (Table 1). Next, we leveraged unique knowledge from various other CF programs to examine knowledge related to IMP3. First, we queried the LINCS L1000 data [86] from the LINCS program [61] converted into RNA-seq-like LINCS L1000 Signatures [87] using the SigCom LINCS API [62] to identify mimickers or reversers small molecules and CRISPR KOs that maximally impact the expression of IMP3 in human cell lines. These potential drugs and targets were filtered using the CF IDG program’s list of understudied proteins [88] to produce a set of additional targets. Next, IMP3 was searched for knowledge provided by the Metabolomics Workbench MetGENE tool [74]. MetGENE aggregates knowledge about pathways, reactions, metabolites, and studies from the Metabolomics Workbench CF supported resource [73]. The Metabolomics Workbench was searched to find associated metabolites linked to IMP3. Furthermore, we leveraged the Linked Data Hub (LDH) API [82] to list knowledge about regulatory elements associated with IMP3. Finally, the GlyGen database [63] was queried to identify relevant sets of proteins that are the product of the IMP3 genes, as well as known post-translational modifications discovered on IMP3. The discovery of IMP3 is not completely novel, IMP3 has been previously reported to be aberrantly expressed in several cancer types and its high expression is associated with poor prognosis [89].

**Table 1.** Ranked list of targets identified by TargetRanger to be highly expressed in the tumor sample and lowly expression across normal tissues from GTEx.

| Gene | Z-score |
| --- | --- |
| IMP3 | inf |
| ARHGDIA | inf |
| GPRIN1 | 7.23 |
| CARM1 | 6.98 |
| JSRP1 | 6.70 |
| SLC7A6 | 6.60 |
| NBPF15 | 5.78 |
| RABGEF1 | 5.76 |
| HPS4 | 5.64 |
| ANKRD39 | 5.21 |

#### Use Case 1: Explaining Drug-Drug Interactions

This workflow takes as input an adverse event term and two drugs. The adverse event is identified in several databases that contain gene sets already associated with the adverse events and mammalian phenotypes related to the adverse event. Namely, matching adverse events and mammalian phenotypes are identified from the GWAS Catalog [90], MGI Mammalian Phenotype ontology [91], and from the Human Phenotype Ontology (HPO) [92]. A set of consensus genes associated with the matching terms is assembled. Then, the workflow queries the LINCS L1000 chemical perturbation signatures [62] with the two input drugs to find gene sets that are consistently up- or down-regulated by the treatment of human cell lines with these drugs. The consensus gene sets impacted by the drugs, and the gene set related to the adverse events are then compared and visualized using a SuperVenn diagram to highlight overlapping genes between these sets. Genes of interest are those affected by both drugs and are associated with the phenotype. Such overlapping genes can be further interrogated individually for evidence in the literature, or as a gene set using enrichment and network analyses.

To demonstrate the workflow for a specific instance, we start with the adverse event “bleeding” and the drugs warfarin and aspirin. It is known that these drugs interact to increase the risk of internal bleeding [93] but the exact intracellular mechanism of such interaction is still not fully understood. The workflow starts with selecting “bleeding” as the search term. Gene sets with set labels containing the word bleeding were queried from Enrichr [1]. Identified matching terms from the GWAS Catalog 2019 [2], MGI Mammalian Phenotype Level 4 2019 [3] and the Human Phenotype Ontology [4] libraries are then assembled into a collection of gene sets. A GMT file is extracted from the Enrichr results for all the identified gene sets from each library and then these are combined using the union set operation. Gene sets with set labels containing the terms warfarin and aspirin were next identified from the LINCS L1000 Chem Pert Consensus Sigs [5] library. The gene sets collected for each drug were combined into one gene set library. The collection of gene sets was then visualized with a SuperVenn diagram (Fig. 5). This analysis identified 243 genes up-regulated and 245 genes down-regulated by warfarin; 249 genes up-regulated and 244 genes down-reguklated by aspirin, 85 genes associated with bleeding from MGI, and 35 from HPO. Only one gene, namely THBS2, is up regulated by both drugs, and is also associated with bleeding related phenotype in MGI. While the gene SLC7A11 is downregulated by both drugs and is linked to an MGI bleeding phenotype. THBS2 is a member of the thrombospondin family, and as such it plays a critical role in coagulation. It was shown that knockout mice of THBS2 have an increased bleeding time phenotype (MP:0005606) [94] and THBS2 is a potent inhibitor of tumor growth and angiogenesis [95]. It is difficult to explain why both drugs are found to up-regulate this gene. The expected effect is that these drugs would reduce the expression of the genes to reduce coagulation. At the same time, both drugs are also found to down-regulate the expression of the amino acid transporter SLC7A11. SLC7A11 knockout mice also have an increased bleeding time phenotype (MP:0005606), and mutations in this gene have been implicated in many acute human diseases through induction of ferroptosis [96,97]. Hence, for SLC7A11 the direction of the impact of the drugs on its expression is consistent with other prior evidence.

**Fig. 5.**
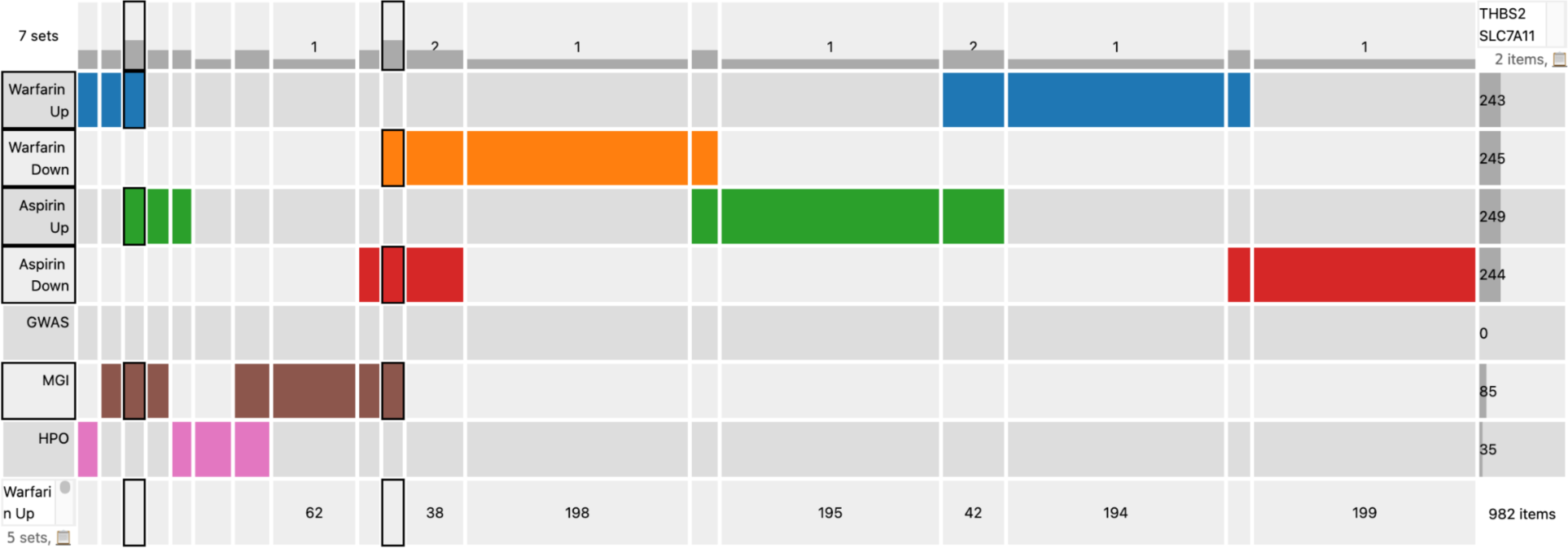
SuperVenn diagram to visualize the overlap between sets of genes that are up and down regulated by aspirin and warfarin based on LINCS L1000 signatures as well as knockout mouse, HPO, and GWAS phenotypes associated with the term “bleeding”. The permanent URL for accessing the workflow in the PWB platform is: https://playbook-workflow-builder.cloud/report/d94b8b0a-81cc-708c-e200-e00ef3451da0

#### Use Case 11: Related Proteins/Metabolites across DCCs

The enzyme ribulose-5-phosphate epimerase (RPE) participates in the catalysis of the interconversion of ribulose-5-phosphate (Ru5P) to xylulose-5-phosphate (Xu5P) in the pentose phosphate pathway. A recent study [98] focused on the biophysical and enzymatic characterization of RPE in several organisms. Interestingly, the study suggested that RPE may play a crucial role in protection against oxidative stress. Towards integrative analysis to further elucidate the roles of RPE in various pathways and mechanisms of human disease, we collected knowledge about PRE from various NIH CF programs and other sources. The collected information about PRE includes: 1) Associated metabolites from the Metabolomics Workbench [73]; 2) Expression across human tissues from GTEx [52]; 3) Small molecules and single gene knockouts that maximally induce the expression of PRE from LINCS [62]; 4) Associated variants from ClinGen via LDH [99]; 5) Protein-protein interactions from STRING [77]; and 6) Regulation of PRE by transcription factors from ChEA3 [100]. In addition, the use case converts PRE into a gene set using the Geneshot API [57]. The Geneshot API returns a set of 100 genes that mostly correlate with PRE based on thousands of human RNA-seq uniformly processed from GEO [101]. Co-expression correlations computed from the data processed by ARCHS4 [53]. The comprehensive approach to find knowledge about a single gene is also applied to the generated gene set with all six resources. The final report provides a mechanistic understanding of how RPE can affect various pathways and functions despite not being involved in the pathways and processes directly.

#### Use Case 27: Identifying Gene Regulatory Relationships between Genes, Regulatory Elements, and Variants

This workflow takes as input one or more genes, regulatory elements, or variants. One may then query for regulatory relations of the selected entity type with other entity types. In one application, we may ask what genomic regions regulate a gene of interest and what evidence supports that regulatory relationship. We start the workflow by providing the gene of interest as input. We first focus on regulatory elements that are in the vicinity of the gene body identified using the epigenomic data from NIH Roadmap Epigenomics [94] and ENCODE projects stored in the ENCODE SCREEN database [83]. Regulatory evidence associated with the SCREEN regulatory elements was connected to genes and variants using CFDE LDH [82], a graph-based database that facilitates the linking of findable, accessible, interoperable, and reusable (FAIR) [102] information about genes, regulatory elements, and variants retrieved through well-documented RESTful APIs. The available regulatory information includes: 1) Variants associated with regulatory elements from the ClinGen Allele Registry [81]; 2) Allele-specific epigenomic signatures, such as DNA methylation, histone modifications, and transcription factor binding, from Roadmap Epigenomics [84] and EN-TEx [85] projects; 3) Quantitative trait loci information from GTEx [52] and other studies; and 4) Regulatory element activity, all presented in a tissue- and cell-type-specific manner. The workflow also provides users with commonly used identifiers for variants that fall within a regulatory element of interest, including those from dbSNP [103], ClinVar [104], and the ClinGen Allele Registry [81].

## Conclusions

Here we describe a new web-based interactive workflow construction platform called the Playbook Workflow Builder (PWB). The workflow engine facilitates user traversal through a network of microservices stored in a knowledge resolution graph (KRG). The metanodes are wrappers to external APIs that are executed on-demand with the inputs of the previous step to produce the outputs for the next. The user-friendly web-based interface enables users to extend, branch, and merge a workflow which is executed while it is constructed. Users can construct workflows manually and via a chatbot interface. Notably, the system provides the means to modify decisions at an earlier stage of a workflow and have the workflow following that point re-evaluated to reflect those changes. This makes any given user session a reusable workflow template.

Besides constructing their own workflows, users can also reuse published workflows created by other users. The published workflows contain detailed descriptions of each step, and this provides the ability to construct reports that resemble research publications. These public workflows can be re-executed by interacting with a chatbot via prompts and inject user data or user decisions into the original published workflow. Once the user makes an adjustment, a new workflow is created and executed by the platform and the results presented as they become ready. The automatic description about the workflow may also be adjusted to reflect the user’s specific changes. This newly modified workflow automatically becomes persistent with a unique citable and publishable URL. Some features of the platform require users to log in, such as for uploading files, saving, and publishing workflows, contributing suggestions, and using several features such as publishing workflows as BioCompute Objects or operating the playbook within CAVATICA’s cloud resources.

So far, most of the metanodes and use cases implemented by the PWB platform are related to systems biology, molecular networks, and the analysis of genes, variants, metabolites, and post-translational modifications. The platform is extendible and could be applied to other areas of biomedical research domains such as structural biology, cheminformatics, genomics, and clinical research. In addition, the PWB platform can be applied in other domains besides biomedical research. The chat interface of the PWB also opens opportunities for applications that may enhance the functionality of chat bots and other bots by executing workflows on demand to produce knowledge and understanding that is deeper that would be achieved by large language models (LLMs) and other currently available state-of-the-art AI models.

## Acknowledgements

This project was funded by NIH grants OT2OD036435 (CFDE Workbench), OT2OD030160 (LINCS DCC), OT2OD030544 (MW DCC), OT2OD030547 (ERCC DCC), OT2OD030162 (KF DCC), and OT2OD032092 (GlyGen DCC),

