## Supporting Figures for "Playbook Workflow Builder: Interactive Construction of Bioinformatics Workflows from a Network of Microservices"

#### Fig. S1

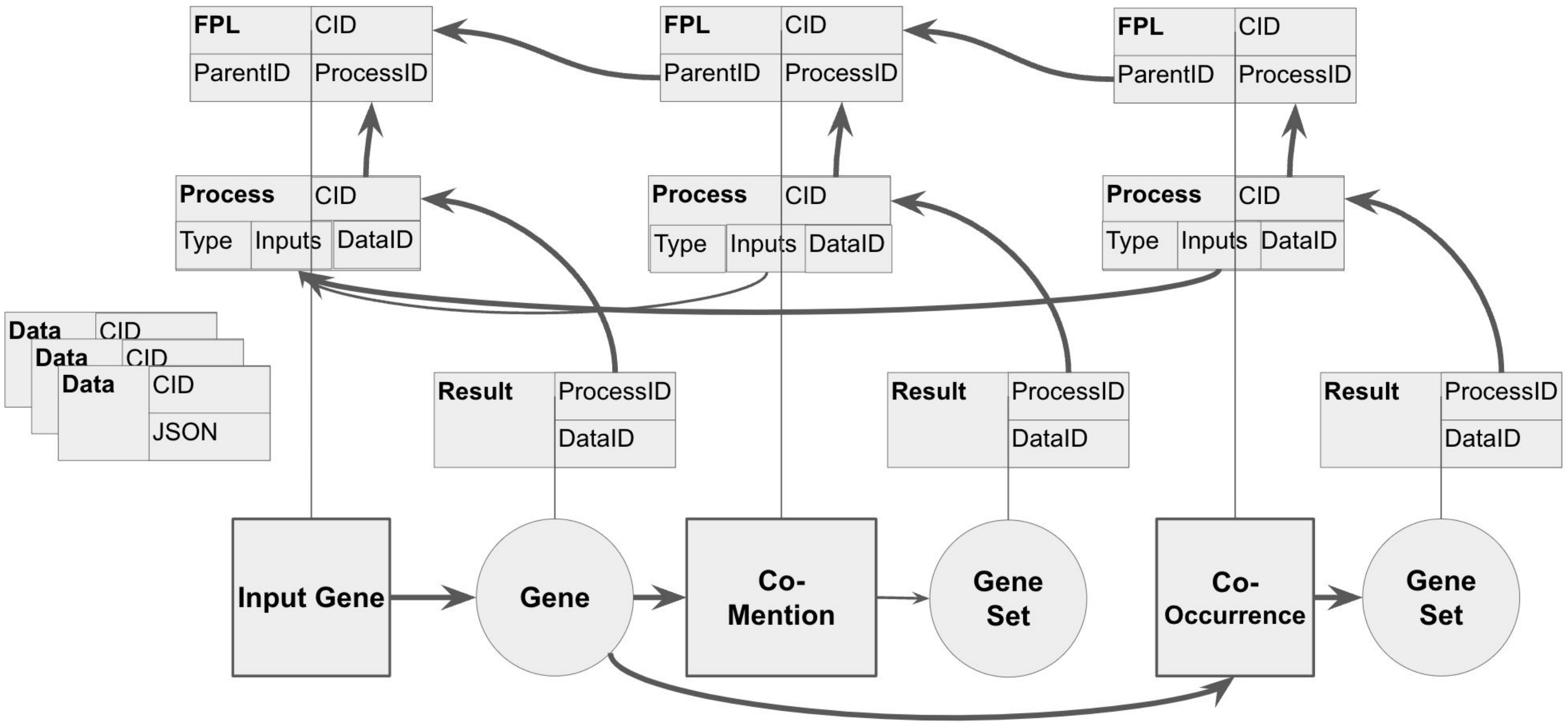
FPL
CID
ParentID
ProcessID
FPL
CID
ParentID
ProcessID
FPL
CID
ParentID
ProcessID
Process
CID
Type
Inputs
DataID
Process
CID
Type
Inputs
DataID
Process
CID
Type
Inputs
DataID
Data
CID
Data
CID
Data
CID
JSON
Result
ProcessID
DataID
Result
ProcessID
DataID
Result
ProcessID
DataID
Input Gene
Gene
Co-Mention
Gene Set
Co-Occurrence
Gene Set

**Fig. S2 A**

### *Use Case 1 - Explain Drug-Drug Interactions*

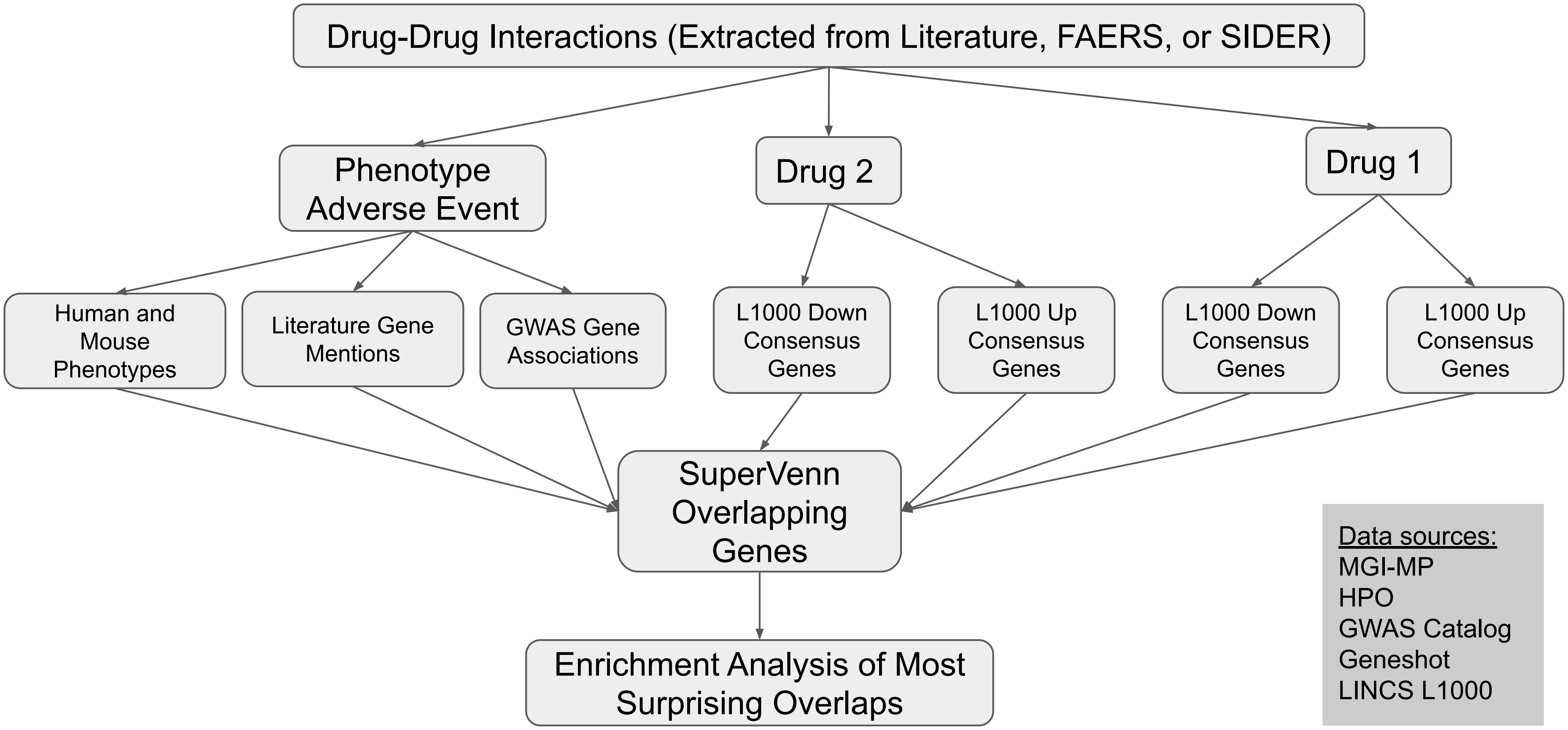

### *Use Case 2 - Explain MOAs of Side Effects for Approved Drugs*

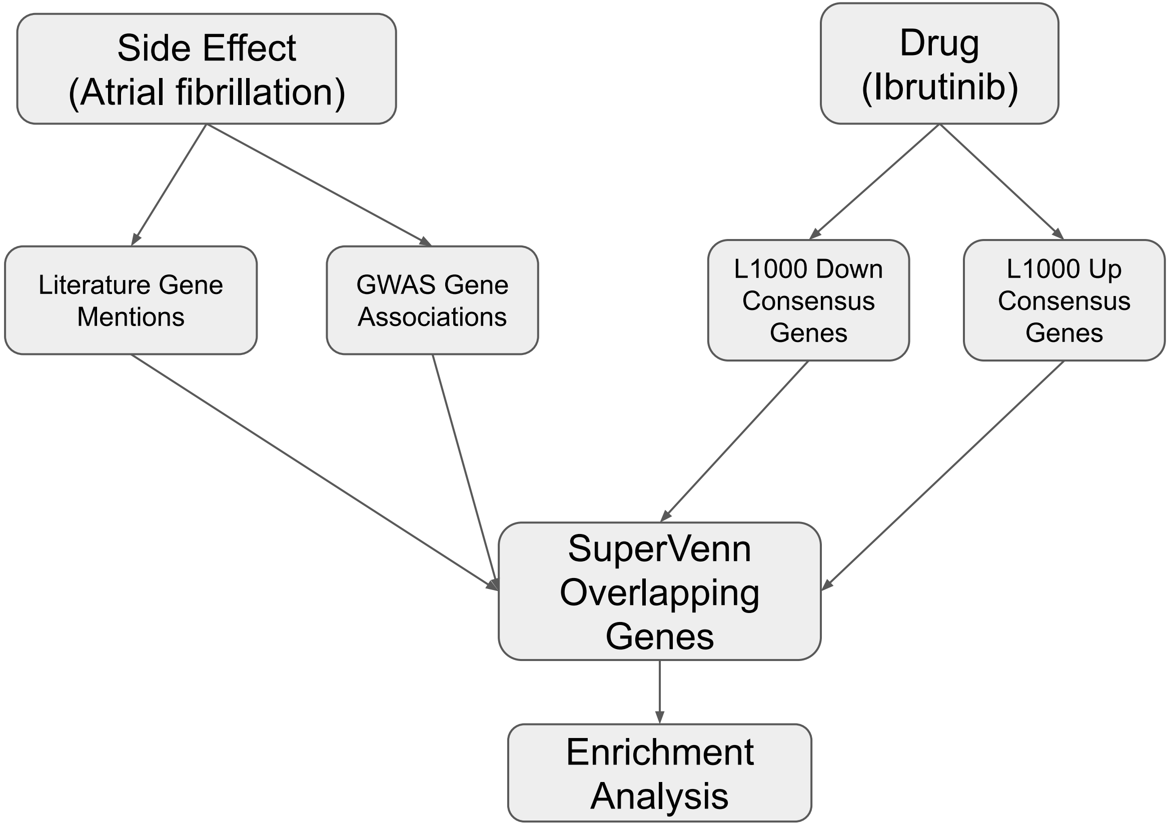


<u>Data sources:</u>
GWAS Catalog
Geneshot
LINCS L1000

**Fig. S2 B**

### *Use Case 3 - Compounds to Reverse Disease Signatures*

<u>Data sources:</u>
GTEx
GEO
LINCS L1000

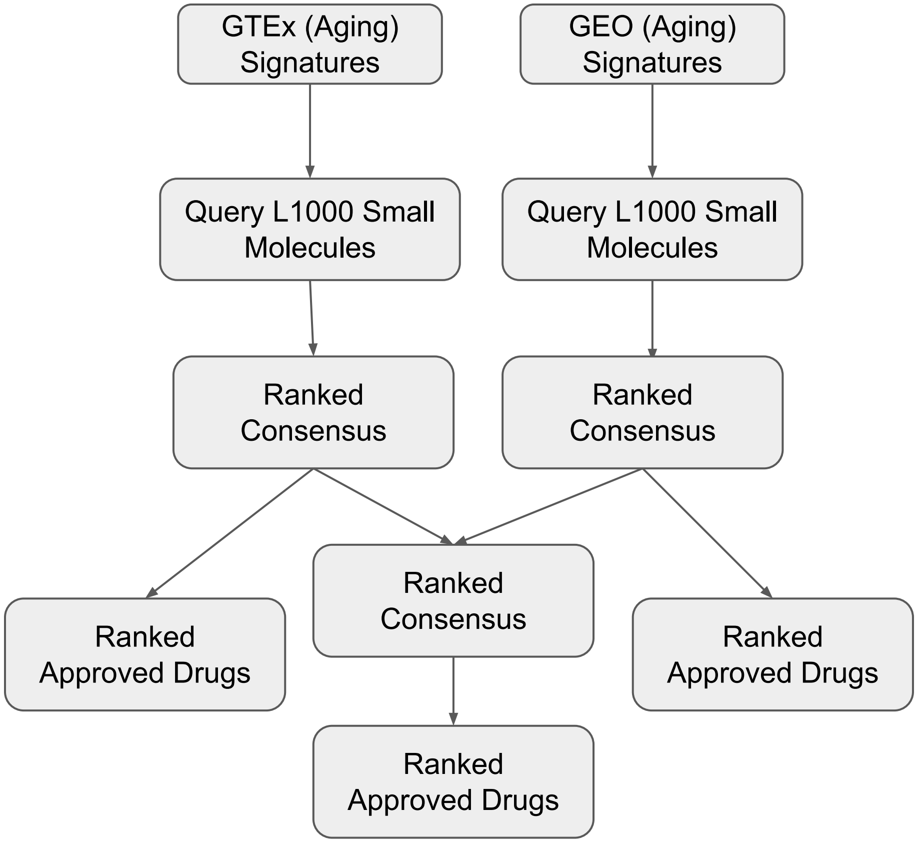


**Fig. S2 C**

**Fig. S2 D**

### *Use Case 4 - KLF4 Targets in GTEx*

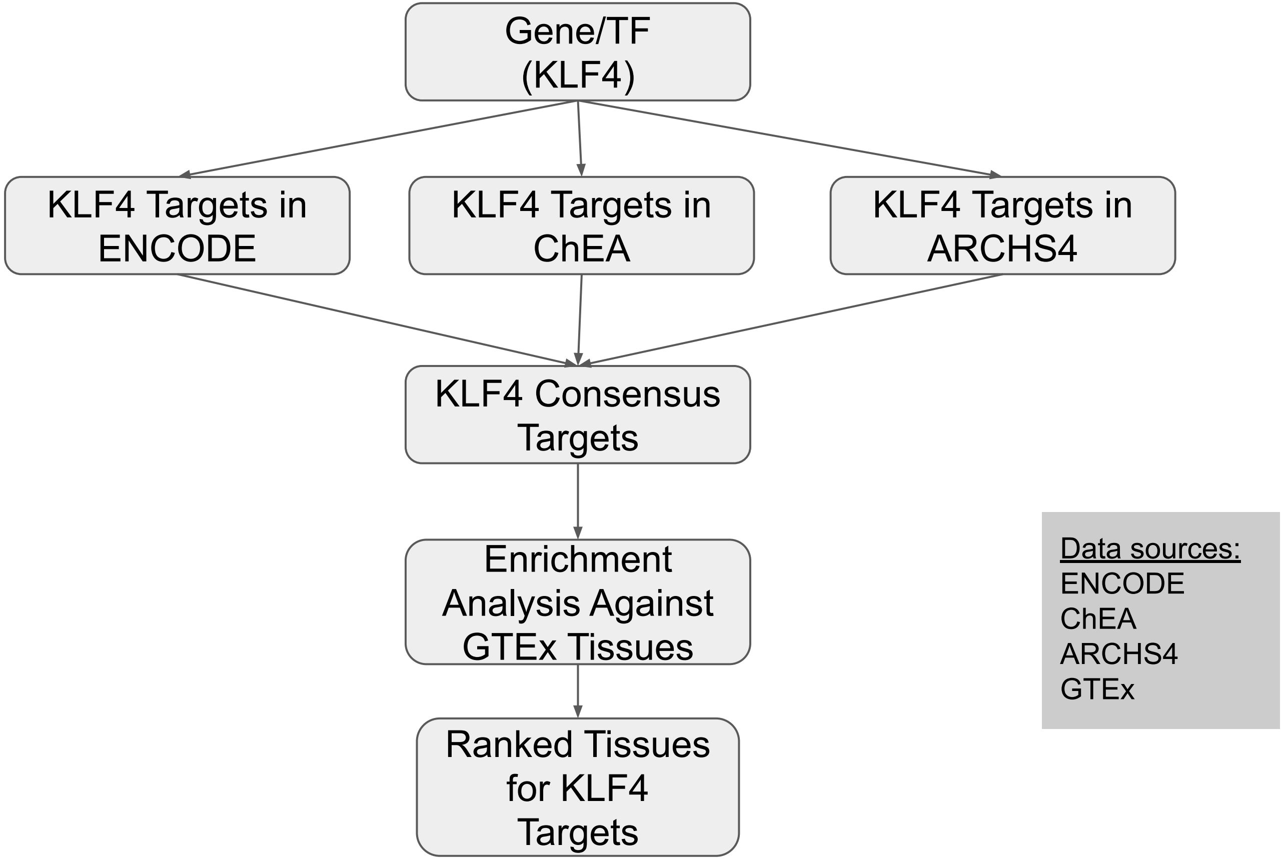

### *Use Case 5 - Small Molecules to Induce a Biological Process (e.g. Autophagy)*

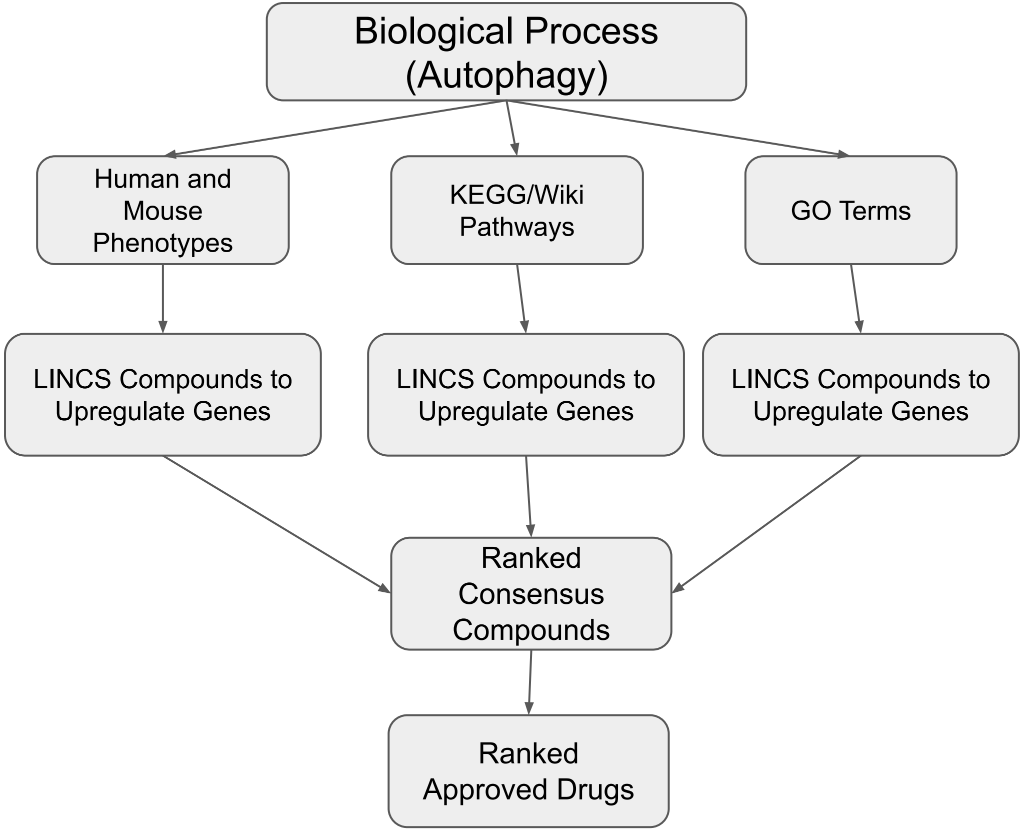


Data sources:
HPO
MGI-MP
KEGG
WikiPathways
GO
LINCS L1000

**Fig. S2 E**

**Fig. S2 F** *Use Case 6 - CFDE Knowledge about a Gene (KLF6)*

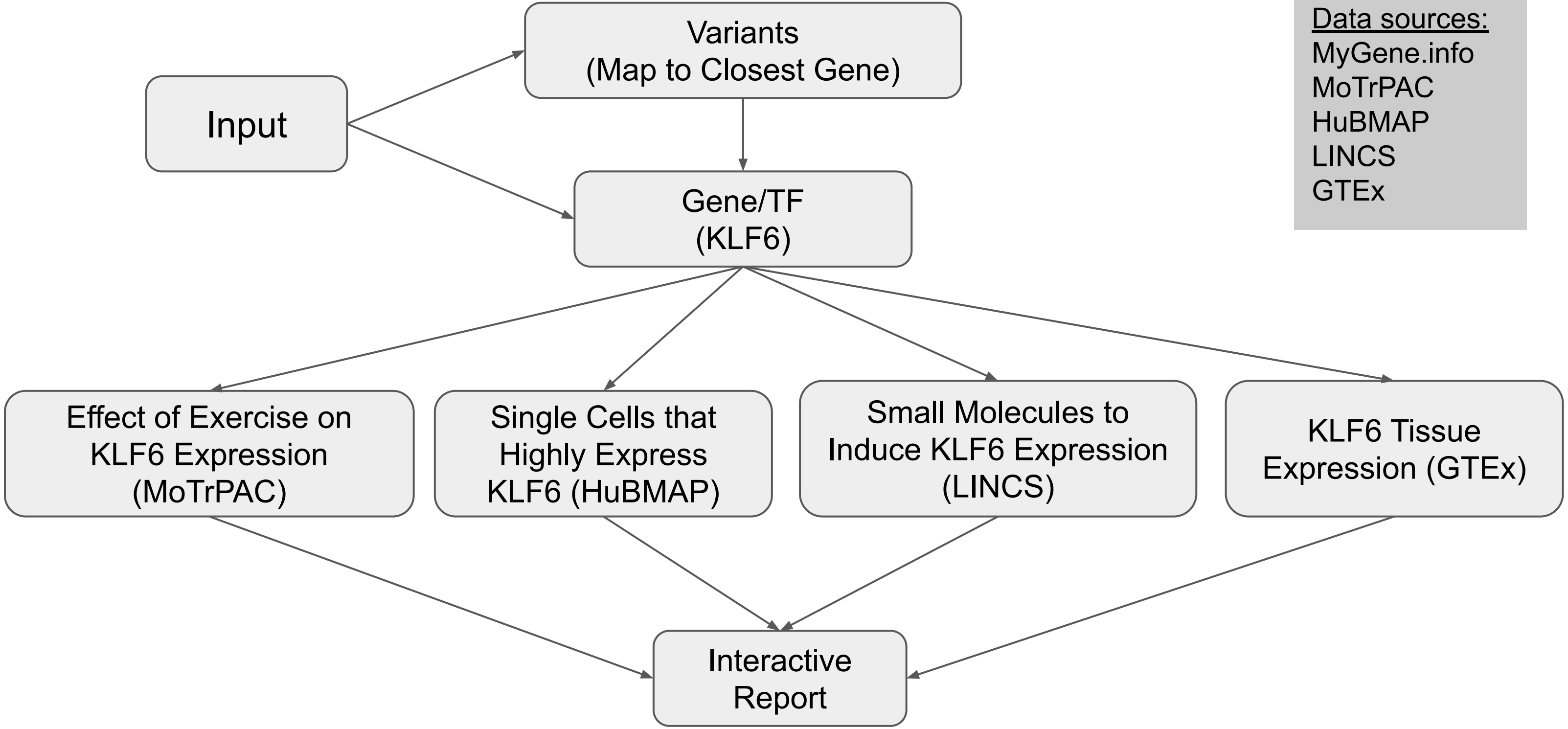


Data sources:
MyGene.info
MoTrPAC
HuBMAP
LINCS
GTEx

**Fig. S2 G**

### *Use Case 7 - Gene/Variant Expression*

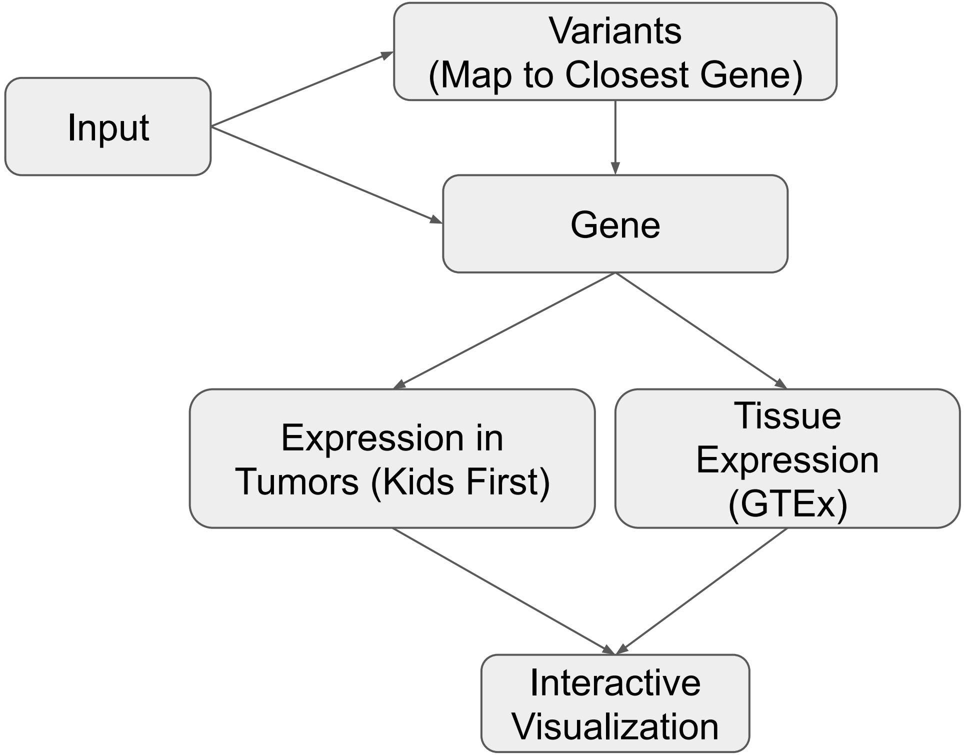


<u>Data sources:</u>
Kids First
GTEx

### *Use Case 8 - Associations between 2 Genes/Variants*

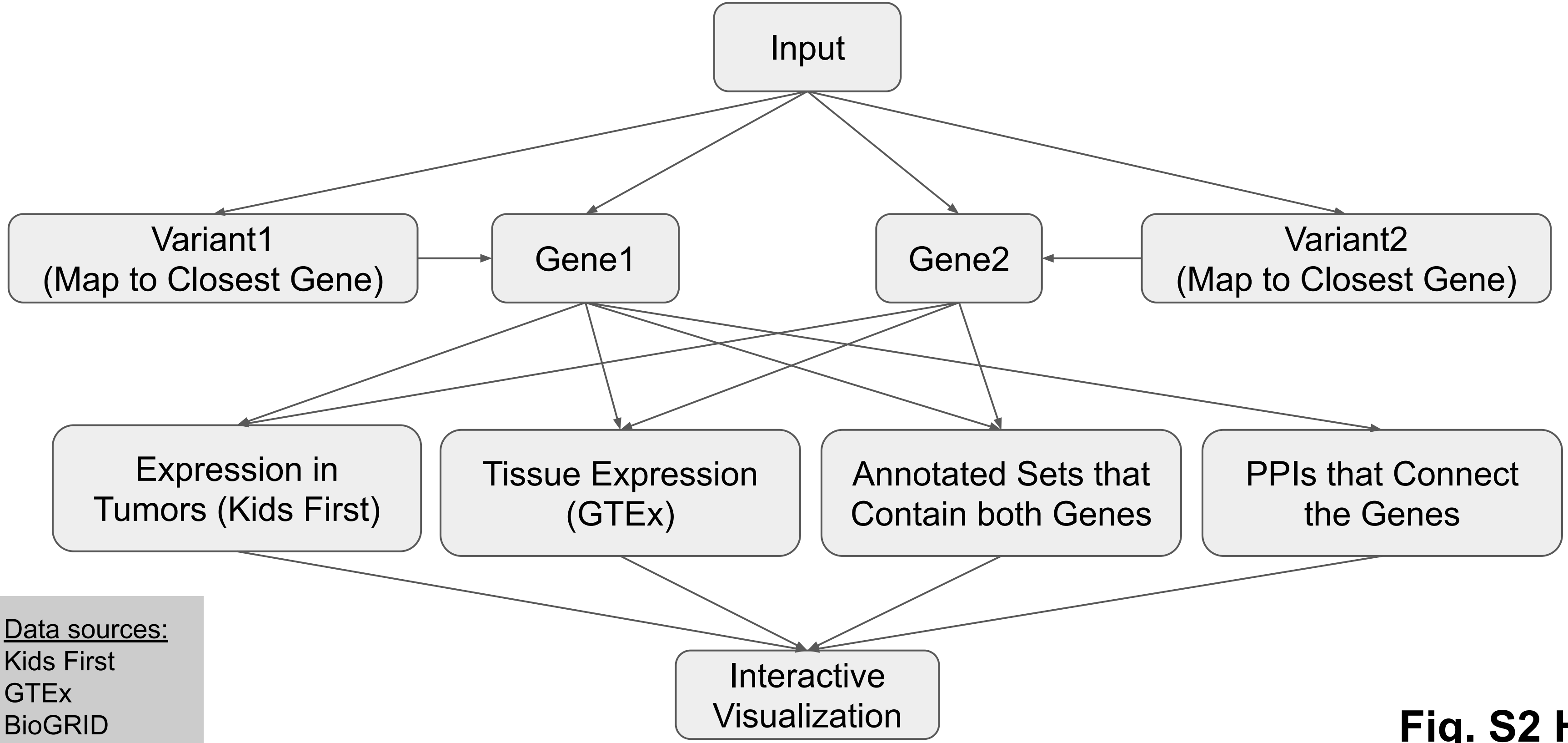


<u>Data sources:</u>
Kids First
GTEx
BioGRID
MyGene.info

**Fig. S2 H**

### *Use Case 9 - Identifying Regulatory Relationships between Genes, Regulatory Regions, and Variants using FAIR Information and Knowledge*

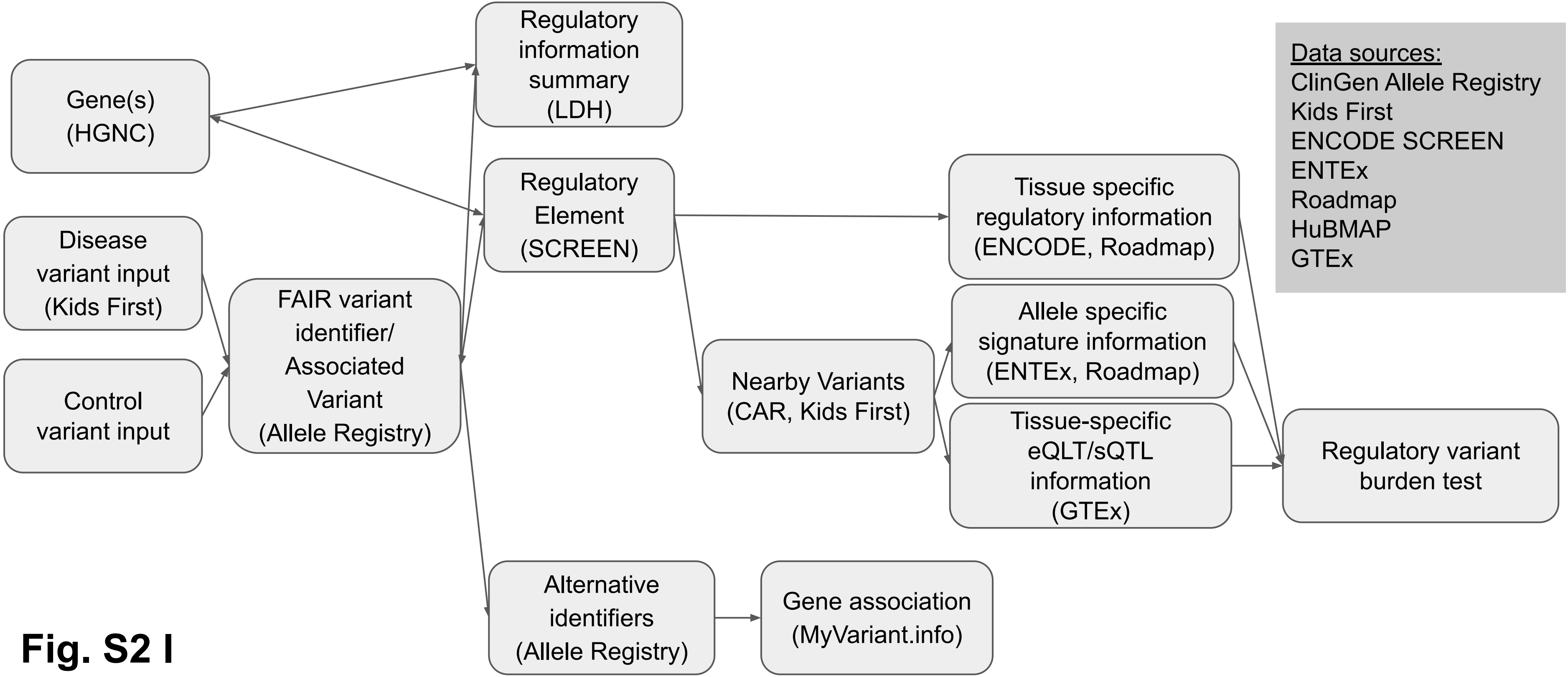


**Fig. S2 I**

*Use Case 12 - SNVs Impact on Glycosylated Sites and Glycogenes Expression*

**Fig. S2 J**

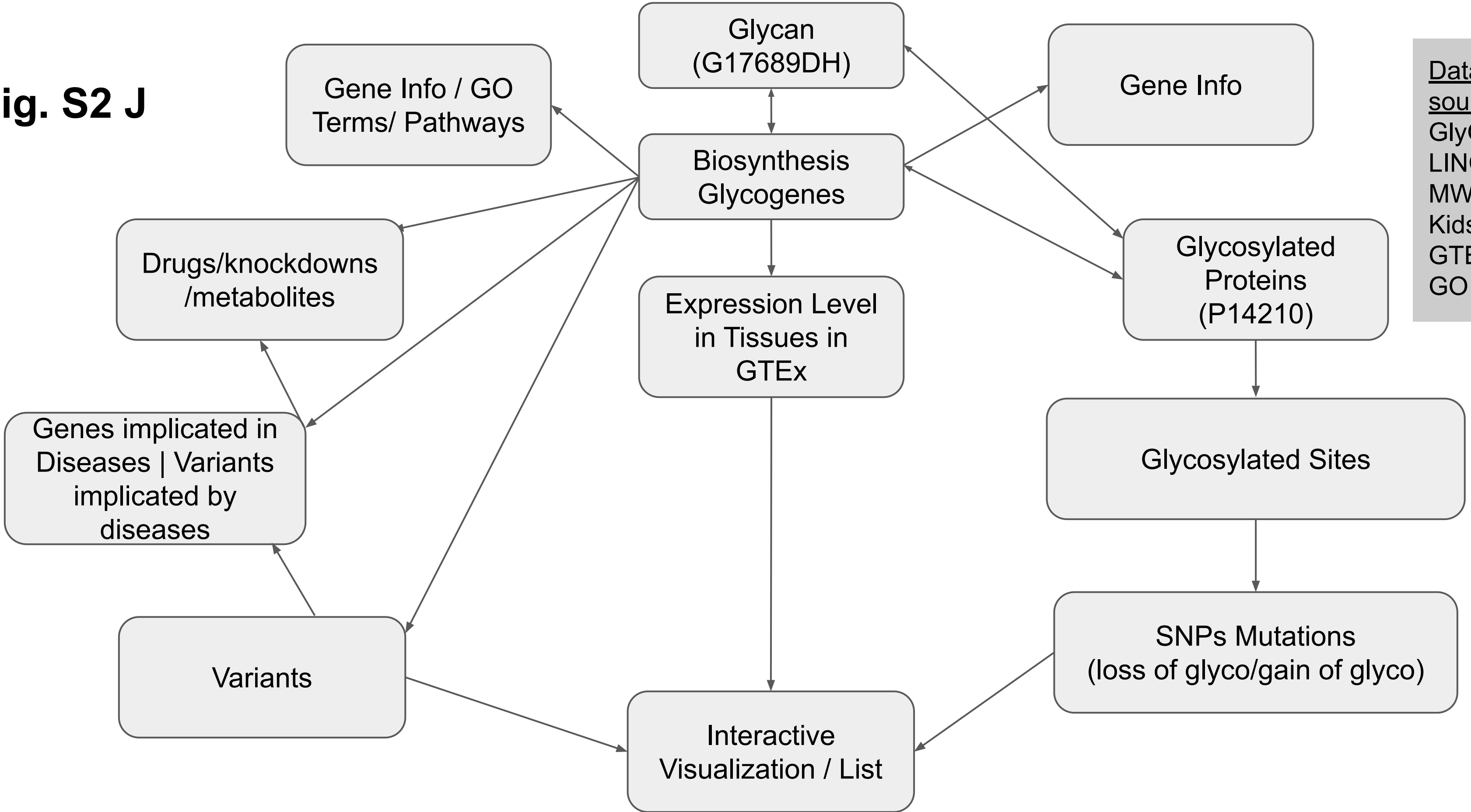


Data sources:
GlyGen
LINCS
MW
Kids First
GTEx
GO

### *Use Case 13 - Prioritizing Targets for Individual Cancer patients*

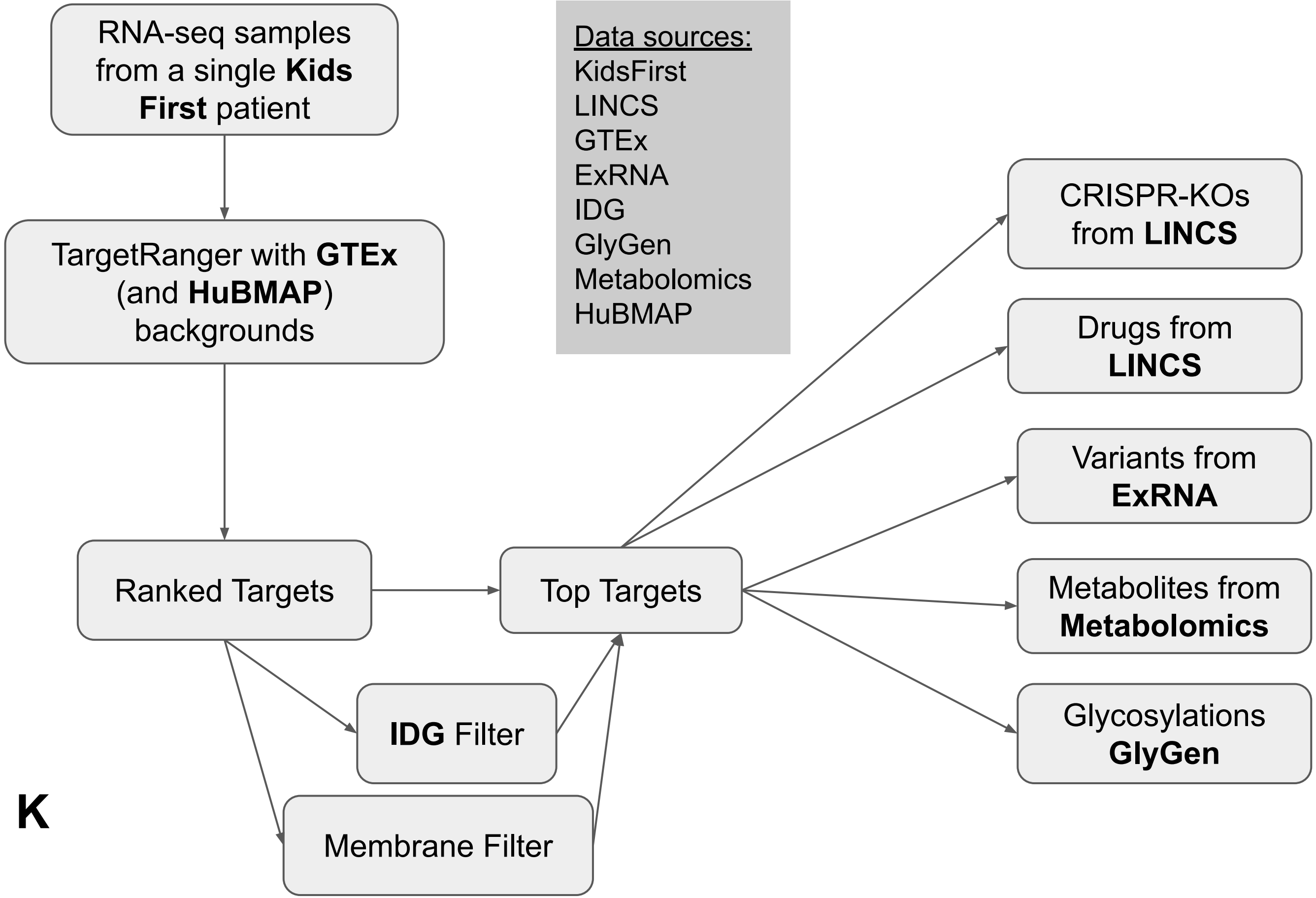


**Fig. S2 K**

### *Use Case 21 - Process a GSE Study and Perform Enrichment Analysis for the DEGs Against CF Gene Set Libraries*

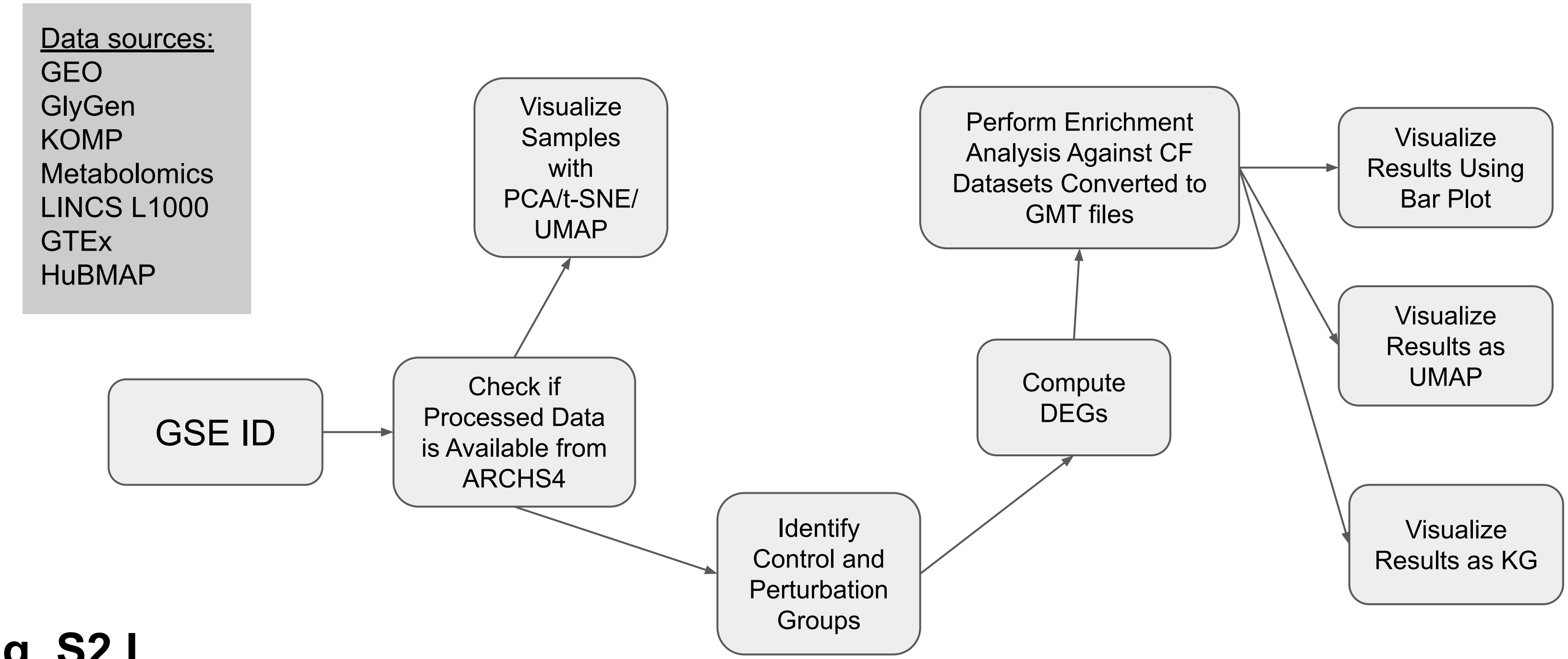


**Fig. S2 L**

### *Use Case 22 - Perform Kinase Enrichment Analysis Followed by Compound Identification from LINCS L1000 and Other Sources*

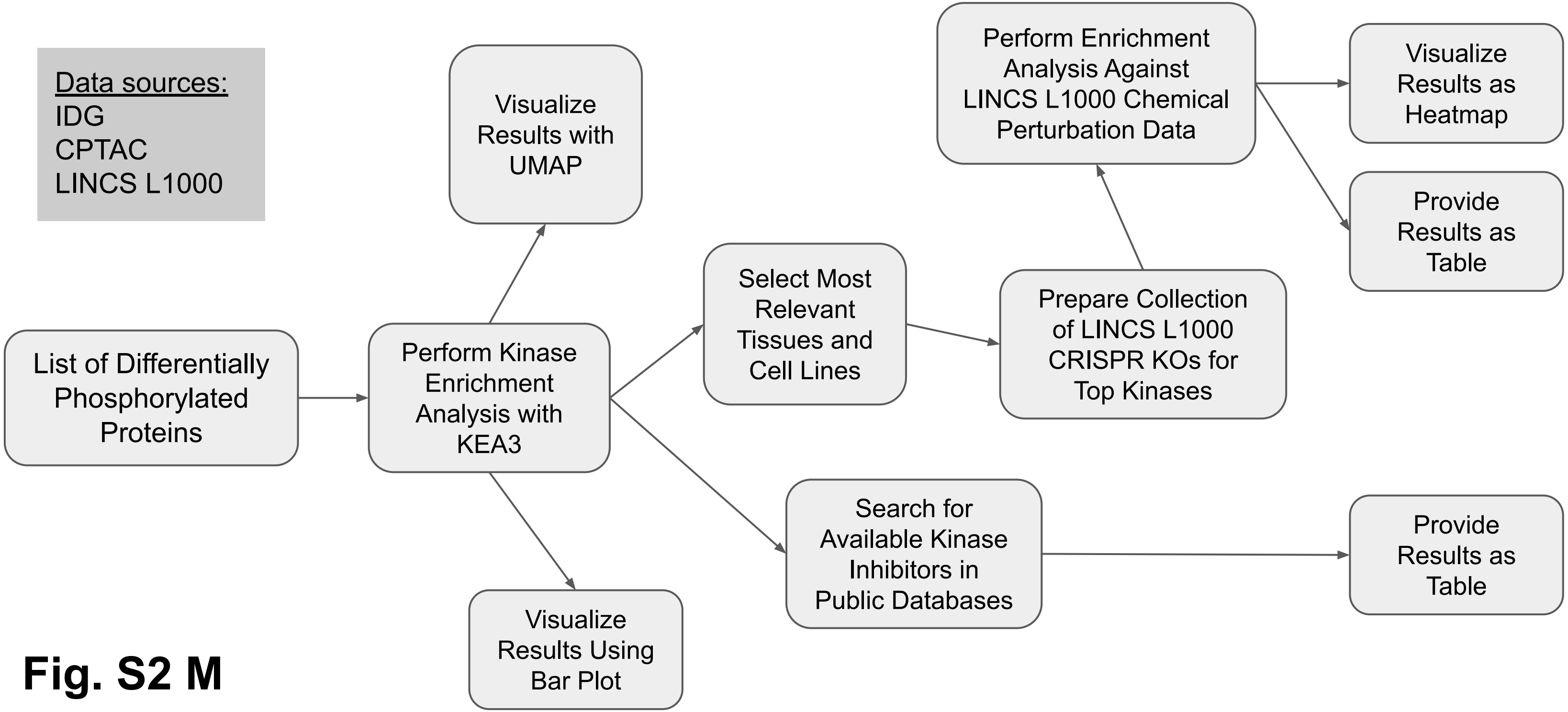


**Fig. S2 M**

### *Use Case 23 - Find Small Molecules and Drugs that Reverse Gene Expression in Young vs. Old Tissues from GTEx (LINCS)*

<u>Data sources:</u>
GTEx
LINCS L1000

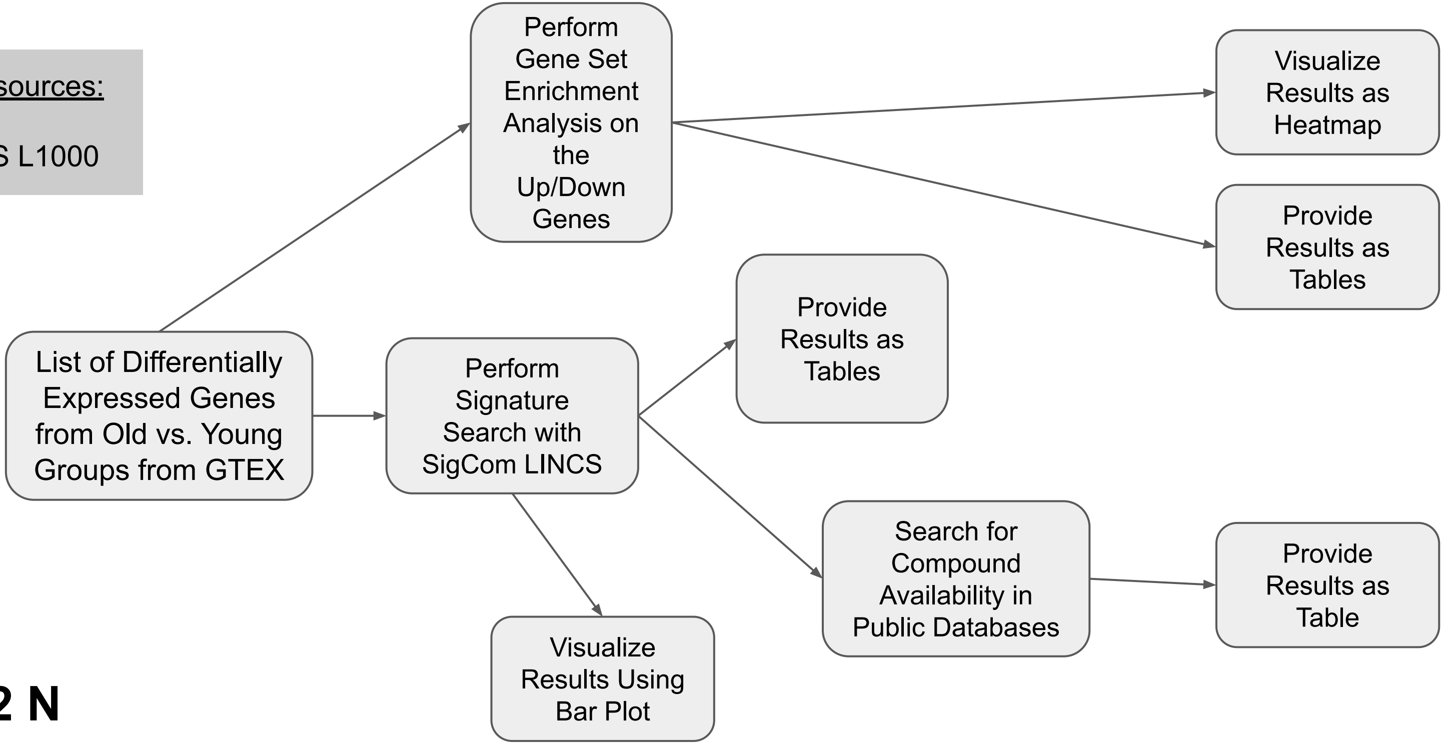


**Fig. S2 N**

### *Use Case 24 - Find Small Molecules and Drugs that Mimic Gene Expression Signature Changes Due to Exercise from MoTrPAC*

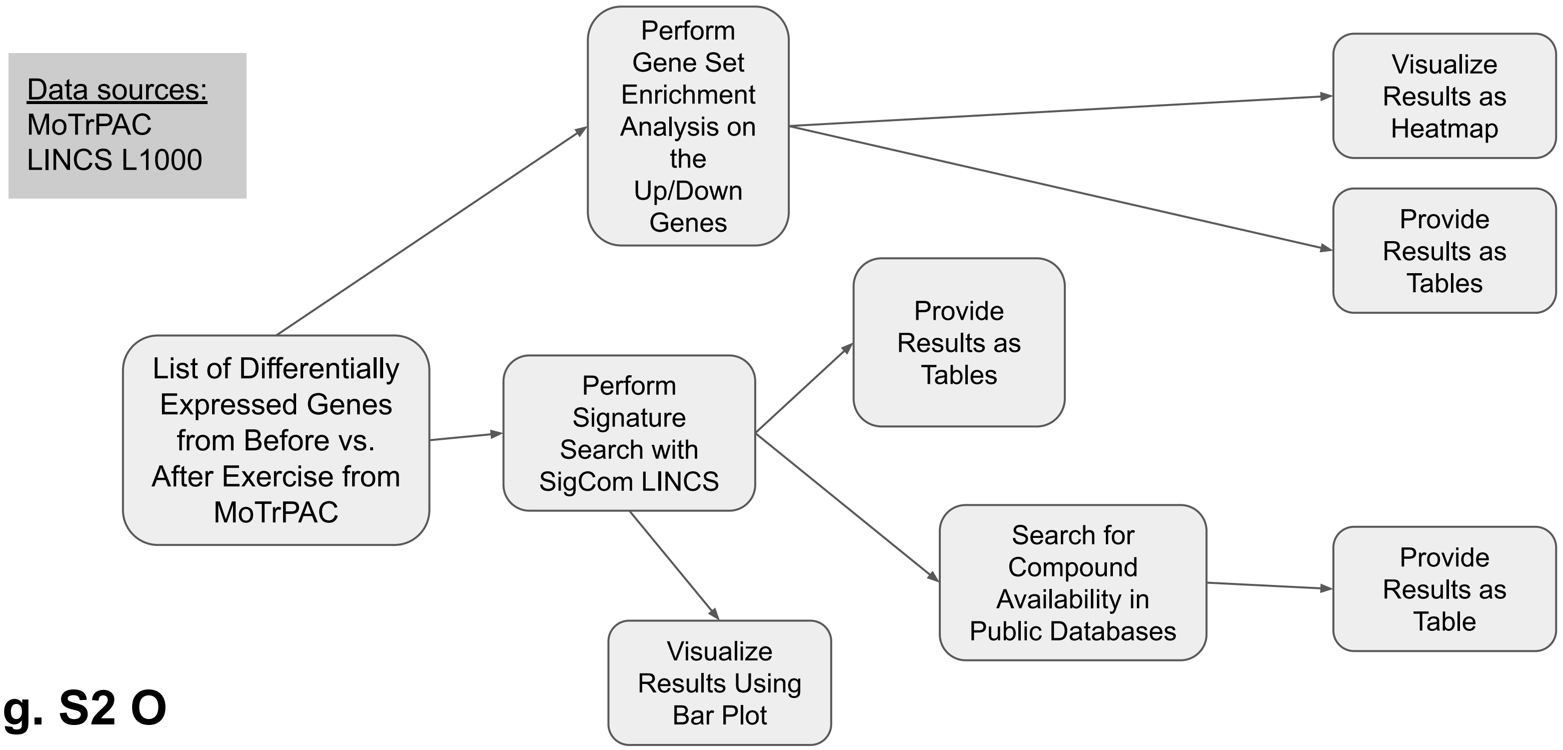


**Fig. S2 O**

### *Use Case 25 - Predict Small Molecules that are Likely to Induce Specific Birth Defects*

<u>Data sources:</u>
MGI-MP
HPO
PubMed-Geneshot
LINCS L1000
GWAS Catalog

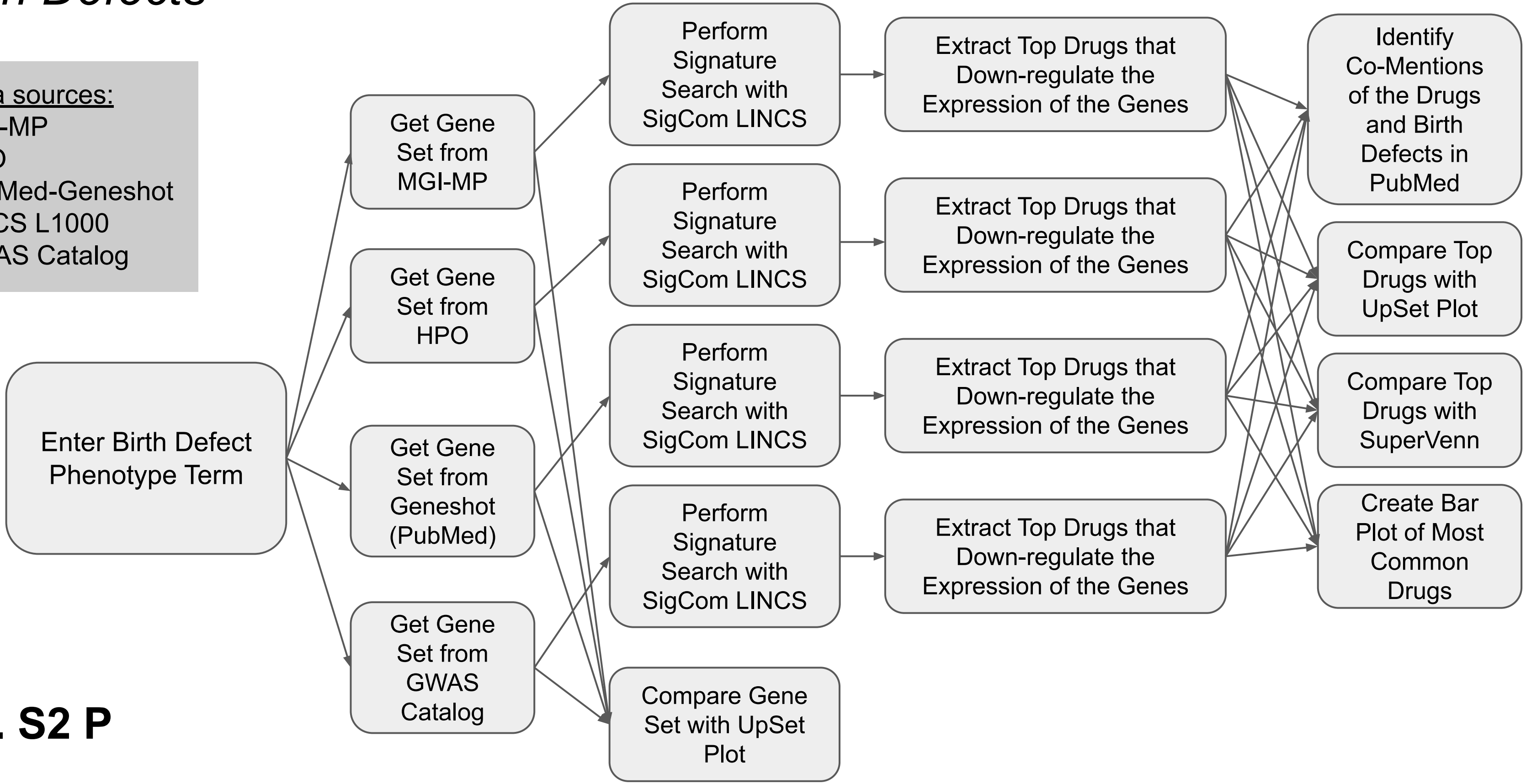


**Fig. S2 P**

### *Use Case 26 – PNPLA3 Related Proteins/Metabolites across DCCs*

Data sources:
LINCS L1000
STRING
ChEA
GTEx
Enrichr
KEGG
GO
MSigDB
MW

**Fig. S2 Q**

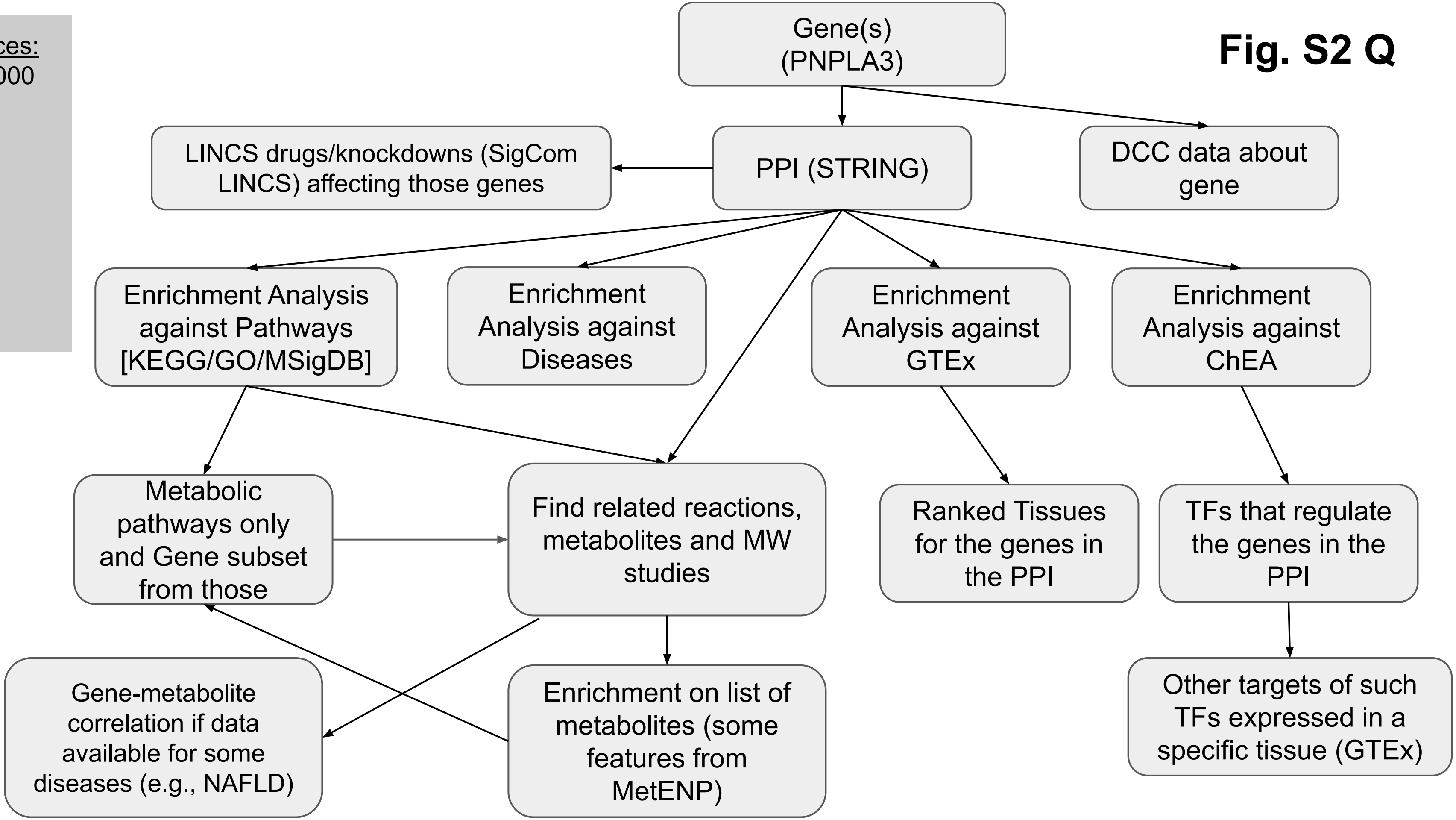

**Fig. S2 R**

### *Use Case 27 - Linking FAIR Regulatory Information for Genes, Regulatory Regions, and Variants*

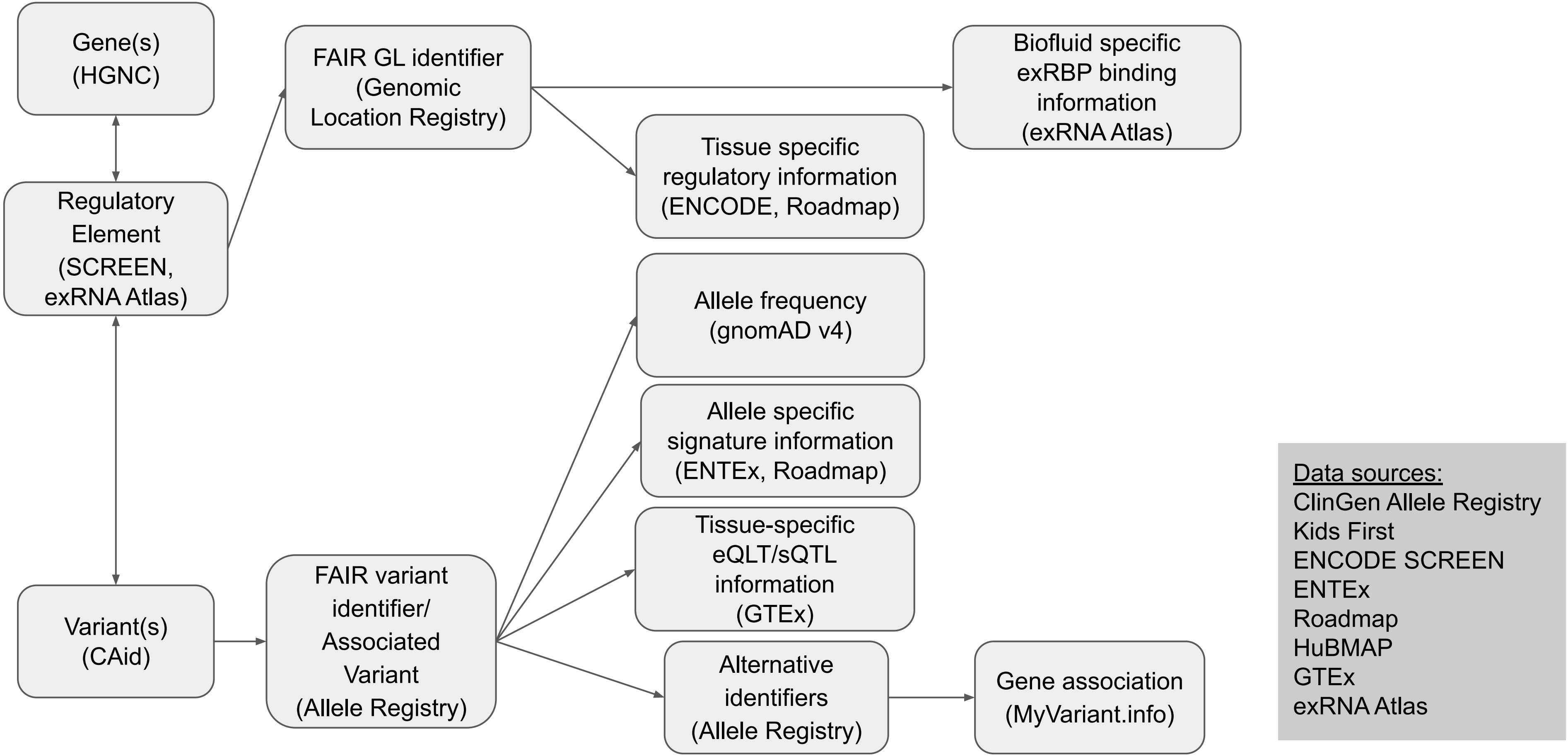

**Fig. S2 S**

### *Use Case 28 - Discovery of De Novo Pathogenic Regulatory Variants from WGS Data*

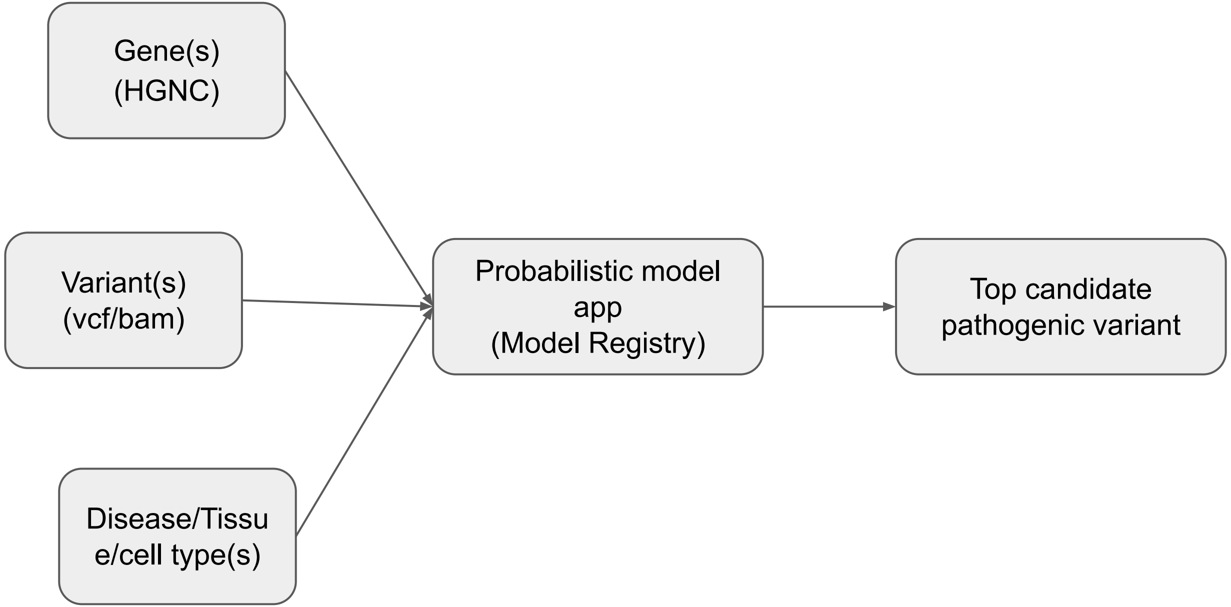


<u>Data sources:</u>
ClinGen Allele Registry
Kids First
ENCODE SCREEN
ENTEx
Roadmap
HuBMAP
GTEx
exRNA Atlas

**Fig. S2 T**

### *Use Case 29 - Form Novel Hypotheses with CFDE Gene Sets and Rummagene*

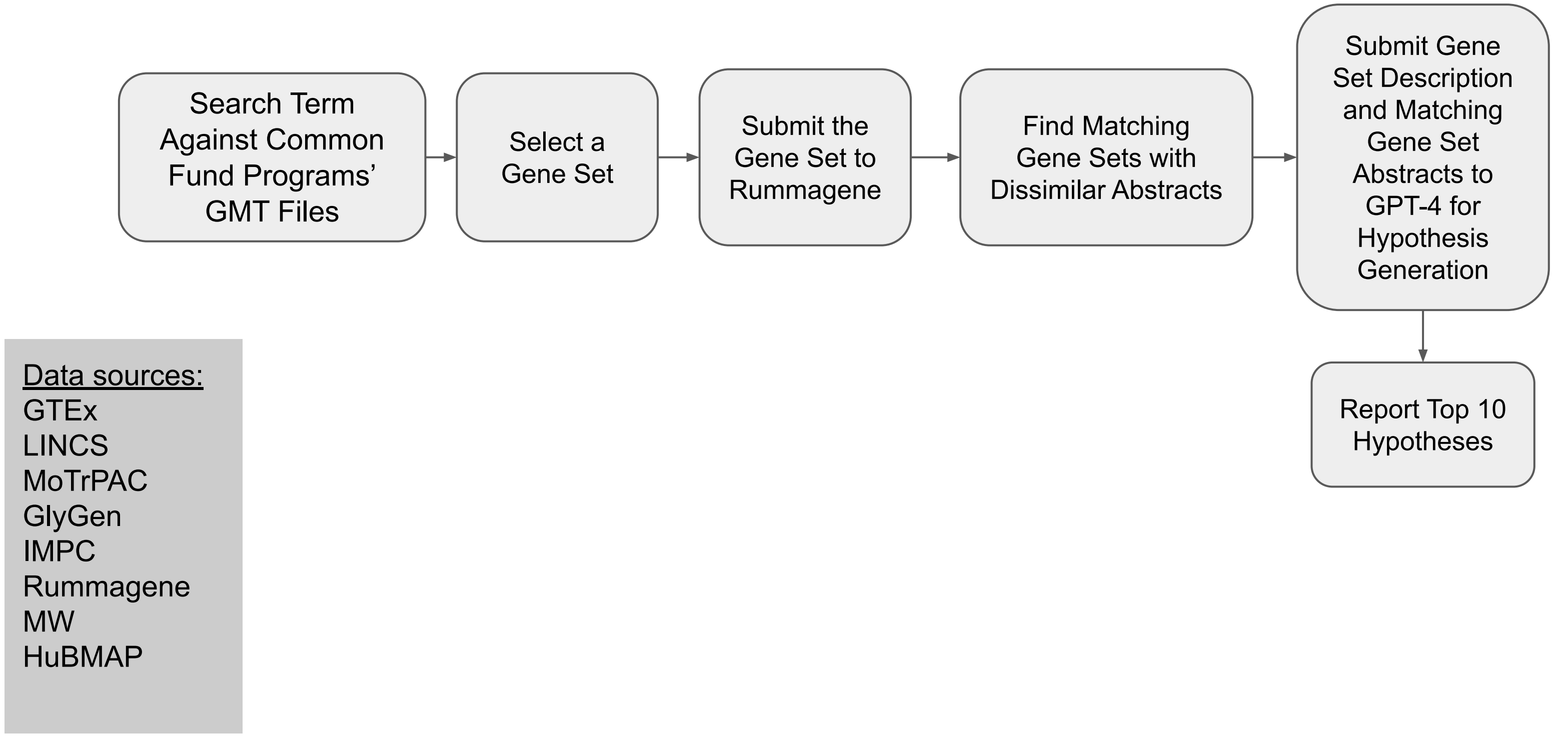

**Fig. S2 U**

### *Use Case 31 - Find Small Molecule Mimickers for Single Gene Knockouts*

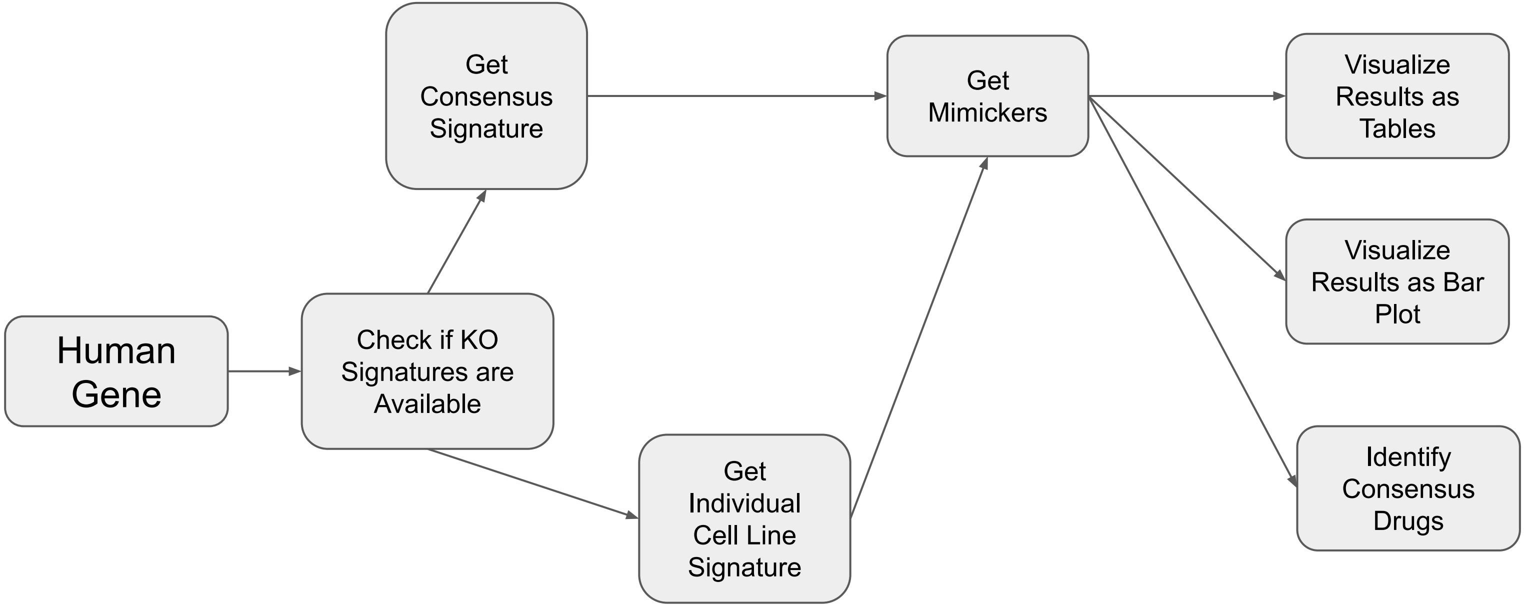


Data sources:
LINCS L1000

**Fig. S2 V**

### *Use Case 32 - Drug Identification for Pediatric Cancer Treatment in Clinical Settings*

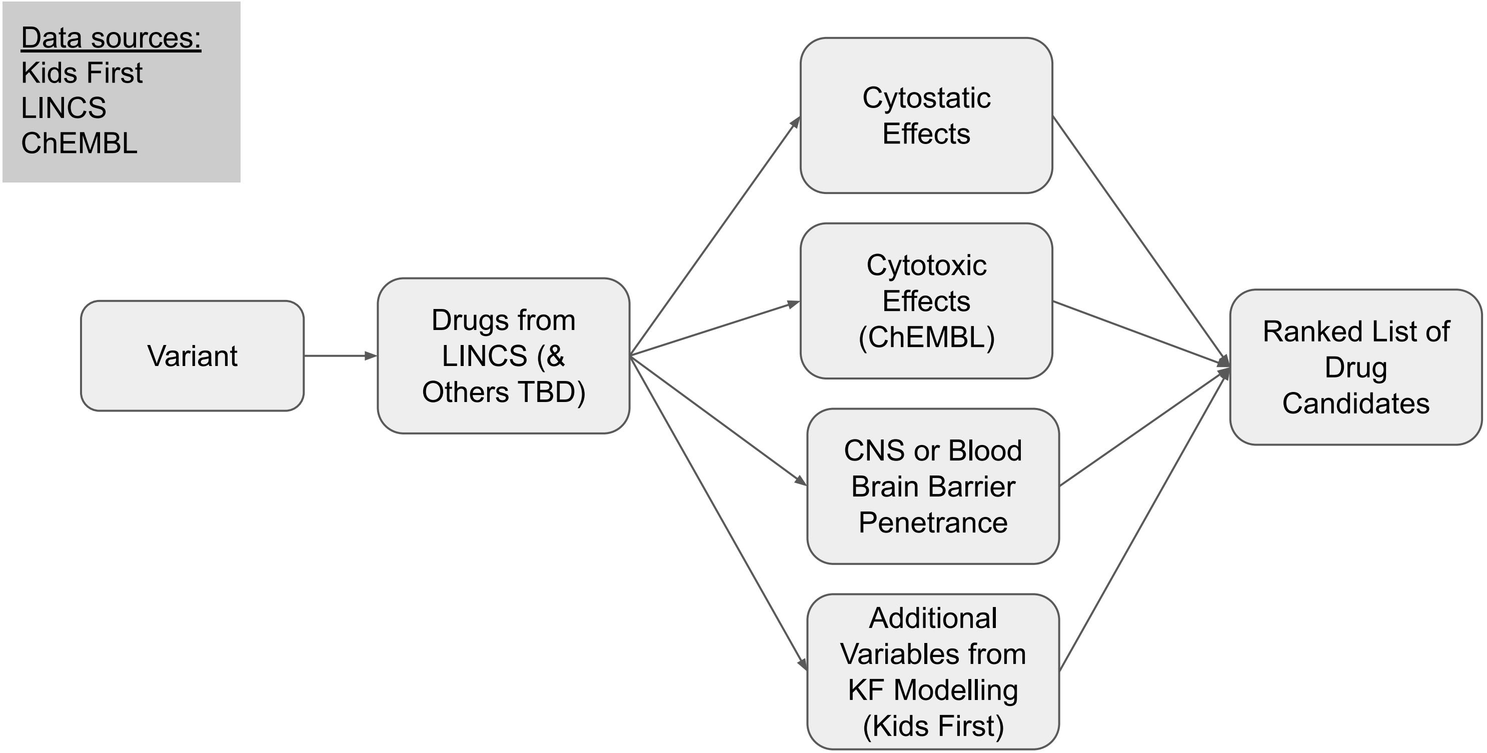
